# A leaky gut dysregulates gene networks in the brain associated with immune activation, oxidative stress, and myelination in a mouse model of colitis

**DOI:** 10.1101/2023.08.10.552488

**Authors:** Jake Sondag Boles, Maeve E. Krueger, Janna E. Jernigan, Cassandra L. Cole, Noelle K. Neighbarger, Oihane Uriarte Huarte, Malú Gámez Tansey

## Abstract

The gut and brain are increasingly linked in human disease, with neuropsychiatric conditions classically attributed to the brain showing an involvement of the intestine and inflammatory bowel diseases (IBDs) displaying an ever-expanding list of neurological comorbidities. To identify molecular systems that underpin this gut-brain connection and thus discover therapeutic targets, experimental models of gut dysfunction must be evaluated for brain effects. In the present study, we examine disturbances along the gut-brain axis in a widely used murine model of colitis, the dextran sodium sulfate (DSS) model, using high-throughput transcriptomics and an unbiased network analysis strategy coupled with standard biochemical outcome measures to achieve a comprehensive approach to identify key disease processes in both colon and brain. We examine the reproducibility of colitis induction with this model and its resulting genetic programs during different phases of disease, finding that DSS-induced colitis is largely reproducible with a few site-specific molecular features. We focus on the circulating immune system as the intermediary between the gut and brain, which exhibits an activation of pro-inflammatory innate immunity during colitis. Our unbiased transcriptomics analysis provides supporting evidence for immune activation in the brain during colitis, suggests that myelination may be a process vulnerable to increased intestinal permeability, and identifies a possible role for oxidative stress and brain oxygenation. Overall, we provide a comprehensive evaluation of multiple systems in a prevalent experimental model of intestinal permeability, which will inform future studies using this model and others, assist in the identification of druggable targets in the gut-brain axis, and contribute to our understanding of the concomitance of intestinal and neuropsychiatric dysfunction.

## 1. Introduction

The intestine is gaining increasing attention as a possible initiator of brain injury in many neuropsychiatric conditions. Emotional dysfunction, cognitive deficits, and corresponding functional connectivity impairments in cortical and limbic areas are well-documented in patients with inflammatory bowel diseases (IBDs), including Crohn’s disease (CD) and ulcerative colitis (UC) (Bernstein et al., 2019; Li et al., 2021; Thomann et al., 2019; Walker et al., 2008). IBD also confers a heightened risk of developing neurodegenerative diseases, including a roughly 50% increased risk of developing multiple sclerosis (MS) and a 20-40% increased risk of developing Parkinson’s disease (PD) (Gupta et al., 2005; Kosmidou et al., 2017; Villumsen et al., 2019; Weimers et al., 2019). Moreover, sub-clinical gastrointestinal dysfunction frequently emerges in neuropsychiatric conditions. For example, patients with autism spectrum disorder (ASD) are seven times as likely to suffer from gastrointestinal symptoms such as diarrhea, constipation, and pain during bowel movements than developmentally intact controls (Chaidez et al., 2014; Thulasi et al., 2019), and constipation is seen in upwards of 60% of PD patients irrespective of IBD (Knudsen et al., 2017). Gut leakiness is of particular concern, potentially admitting injurious microbes from the intestinal lumen to invade tissues, and evidence at functional and molecular levels for intestinal permeability has been found in ASD (Esnafoglu et al., 2017; Fiorentino et al., 2016; Ming et al., 2012) and PD (Aho et al., 2021; Forsyth et al., 2011; Schwiertz et al., 2018). Mechanistic investigation is underway to unearth the cellular and molecular actors underlying the gut-brain connection in these diseased contexts.

Currently, research focuses on the immune system as the injurious mediator between the gut and brain for several reasons. First, many genetic loci whose mutations confer risk to IBD are implicated in the immune system, including *NOD2* (Hampe et al., 2001; Ogura et al., 2001), *LRRK2* (Liu et al., 2015; Rivas et al., 2018), several loci in the *HLA* region (Arimura et al., 2014; Hamza et al., 2010; Stokkers et al., 1999), and several *TRIM* genes including *TRIM10* (Witoelar et al., 2017). Notably, these loci are shared with PD, suggesting that the connection between IBD and PD is immunological (Hui et al., 2018; Kang et al., 2023; Witoelar et al., 2017). Similarly, a large genome-wide pleiotropic study identified a collection of immune system genes enriched associated with cell adhesion, T helper cell differentiation, and Toll-like receptor signaling in the genetic overlaps between gastrointestinal diseases and psychiatric disorders (Gong et al., 2023). Secondly, anti-inflammatory treatments used to combat IBD are protective against neuropsychiatric disease onset. Anti-tumor necrosis factor (TNF) therapy during IBD lowers the later incidence of PD by almost 80% (Peter et al., 2018) and appears to rescue depressive and anxiety symptoms associated with IBD (Gray et al., 2018; Siebenhüner et al., 2021). Finally, several reports highlight positive associations between markers of systemic inflammation, including circulating interleukin (IL-) 6 and C-reactive protein (CRP), or markers of microbial invasion including lipopolysaccharide-binding protein (LBP) with psychiatric symptoms in IBD (Abautret-Daly et al., 2017; Iordache et al., 2022). Despite the clear immunological link between the gut and brain, further studies, including pre-clinical studies that can illuminate cause and effect, are needed to define the exact immunological cascade that drives a gut-to-brain propagation of injury.

The dextran sulfate sodium (DSS) model is perhaps the most frequently employed method used in pre-clinical studies to induce experimental colitis and investigate effects on the gut-brain axis in disease-relevant contexts, especially with a focus on immunity. DSS induces gut leakiness via selective toxicity to intestinal epithelial cells, allowing mucosal immune cells to sense microbial antigens, which leads to a rapid and severe form of colitis (Chassaing et al., 2014; Laroui et al., 2012). The DSS model demonstrates strong translational validity, triggering intestinal immune processes and pathology that mirror those in human IBD (Czarnewski et al., 2019; Holgersen et al., 2015). However, a substantial point of concern in the field is the large degree of variability in generating a robust phenotype representative of disease processes in this model. Specifically, researchers develop their DSS dosing strategy to optimize the phenotype, which may vary significantly between different studies, and research groups generally use different production lots and manufacturers for their DSS reagent, which is a known reproducibility threat (Chassaing et al., 2014; Nell et al., 2010). Additionally, different animal facilities may have differing pathogen loads, which may in turn yield disparate immune responses in the intestine (Cerf-Bensussan and Gaboriau-Routhiau, 2010). Finally, different regions of the colon may be differentially susceptible to DSS-induced colitis, thereby yielding different results in outcome assays. In this study, we examined each of these factors by performing a time-series experiment to investigate the kinetics of DSS-induced damage and inflammatory responses, capturing injury and repair phases and evaluating transcriptomic changes in the gut. We evaluated both distal and proximal colon, the former of which is known to be more vulnerable than the latter (Randhawa et al., 2014). Further, we compared our transcriptomic profiles with those of three other publicly available datasets in which the DSS model was used in a similar fashion to establish universal and experiment-dependent molecular processes in the intestine. As the gut is a source of damage for the brain in this model, we must ascertain whether colitis features are stable across studies and thus whether we can expect that brain consequences of colitis are generalizable.

Crucially, findings from the DSS model support the role of the immune system in a gut-to-brain propagation of injury, with many studies demonstrating inflammation-mediated effects on behavior and brain health in rodents with DSS-induced colitis (reviewed in (Craig et al., 2022)). However, each study published to date focuses on different immunological processes, ignoring other immune processes or non-immune mechanisms altogether, and none identify the cells that may be responsible for brain effects. Additionally, studies provide conflicting results concerning the importance of a systemic immune response (Jackson et al., 2022; Maltz et al., 2022; Vicentini et al., 2022). Here, we adopted an unbiased transcriptomics approach to evaluate perturbances in several brain regions during and after DSS-induced colitis. To provide a comprehensive appraisal of this model, we associated our brain effects with colitis severity indexes, intestinal transcriptomic signatures, and several measures of systemic inflammation. Based on the literature, we hypothesize that DSS induces the infiltration of immune cells into the brain through the activation of systemic inflammation, which triggers persistent neuroinflammation (Houser and Tansey, 2017). We highlight a robust colitis-induced immune activation in the brain along with novel disease processes revealed by our transcriptomic approach, including the dysregulation of myelination processes, oxidative stress, and an imbalanced response to oxygen levels.

## 2. Methods

### 2.1 Mice

C57Bl6/J mice were purchased from Jackson Laboratories, bred to congenicity, and housed until 2-3 months of age prior to experiments while maintained on a 12:12 light-dark cycle with *ad libitum* access to a standard rodent diet chow. All mice were housed in the McKnight Brain Institute vivarium at the University of Florida. All procedures were approved by the University of Florida Institutional Animal Care and Use Committee and followed the *Guide for the Care and Use of Laboratory Animals* from the National Institutes of Health. Mice also received *ad libitum* access to water, which contained dextran sulfate sodium (DSS) depending on group allocation and experimental design, further detailed below.

### 2.2. Cohort 1: Kinetics of DSS-induced gut and brain injury

DSS (2.5%, w/v) was dissolved in autoclaved tap water and placed in sterile water bottles for administration in mouse cages. Six different groups of mice (n = 8-10) received a dosing schedule of varied lengths, according to Figure 1A. The treatment groups were as follows: 5d of DSS, 7d of DSS, 7d of DSS plus 2d of recovery with untainted water, 7d of DSS plus 5d of recovery, 7d of DSS plus 7d of recovery, and 7d of DSS plus 14d of recovery. During DSS treatment, mice were weighed daily and a disease activity index (DAI) was scored as previously described (Supplemental Table 1) (Chassaing et al., 2014). An untreated group (n = 16) was used as controls. After their respective treatment schedules, mice were euthanized via rapid decapitation. Trunk blood was processed to isolate plasma and snap-frozen in liquid nitrogen. Brains were rapidly extracted and bisected, with one hemisphere frozen whole in isopentane and stored at -80°C and the other dissected into cortical, hippocampal, striatal, and ventral mesencephalic tissue which was snap-frozen in liquid nitrogen. Spleens were extracted and weighed as an index of systemic inflammation. Colons were extracted, flushed with PBS to remove fecal matter, and the weights and lengths were recorded as an index of colitis severity. 5mm pieces of the distal and proximal ends were snap-frozen in liquid nitrogen.

**Figure 1:**
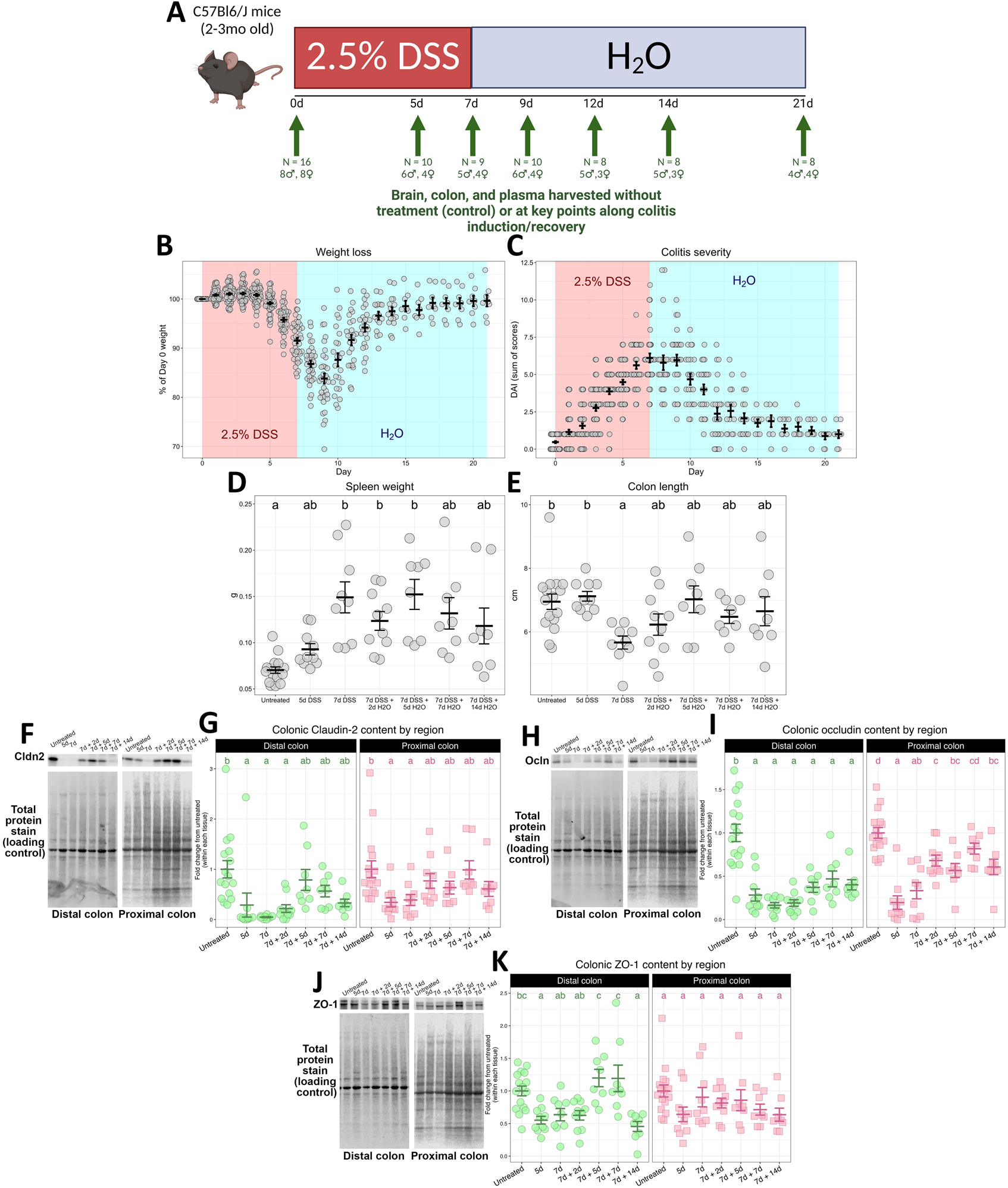
Experimental timeline and key colitis outcomes. **A** Experimental timeline with post-mortem tissue allocation described. **B** Weight loss of mice on DSS, presented as a percentage of their starting weight. **C** Disease activity index (DAI) of mice, shown as the sum of stool consistency scores, fecal occult blood scores, and weight loss scores. **D** Raw spleen weight in grams (one-way ANOVA: Welch’s *F*[6, 22.843] = 10.955, *p* < 0.001). **E** Raw colon length in centimeters (one-way ANOVA: *F*[6, 62] = 2.983, *p* = 0.013). **F** Representative immunoblots for claudin-2 in both colon segments, quantified in **G** (two-way ANOVA: main effect of group – *F*[6, 62] = 7.563, *p* < 0.001; main effect of tissue – *F*[1, 62] = 5.744, *p* = 0.020; interaction - *F*[6, 62] = 1.188, *p* = 0.325). **H** Representative immunoblots for occluding in both colon segments, quantified in **I** (two-way ANOVA: main effect of group – *F*[6, 62] = 6.21.831, *p* < 0.001; main effect of tissue – *F*[1, 62] = 30.032, *p* < 0.001; interaction: *F*[6, 62] = 3.014, *p* < 0.001). **J** Representative immunoblots for ZO-1 in both colon segments, quantified in **K** (two-way ANOVA: main effect of group – *F*[6, 62] = 6.532, *p* < 0.001; main effect of tissue – *F*[1, 62] = 0.049, *p* = 0.826; interaction – *F*[6, 62] = 3.014, *p* = 0.012). For **D**, **E**, **G**, **I**, and **K**, groups that share a letter are not statistically significantly different (*p* > 0.05) according to *post hoc* testing. All summary statistics plotted are mean ±LSEM.

### 2.3. Cohort 2: Immunophenotyping the brain and periphery

A second cohort of mice was treated for 7d of 2.5% (w/v) DSS plus 2d of recovery with untainted water or only untainted water for the entire 9d as a control (n = 12-15/group). Mice were weighed daily, and disease activity index was scored as described on the last day only. After the treatment schedule, mice were euthanized via rapid decapitation. Trunk blood was collected in EDTA-containing vacutainer tubes, and 200µL of whole blood was used to isolate peripheral immune cells. Red blood cells were lysed with a buffer containing 0.15M NH_4_Cl, 10mM KHCO_3_, and 0.1mM EDTA, and cells were washed once in Hank’s Balanced Salt Solution (ThermoFisher, #14175103). Cells were centrifuged at 350*g* for 5min at 4°C and resuspended in a phosphate-buffered saline (PBS) ahead of surface staining for flow cytometry. Brains were rapidly removed and processed for immune cell isolation and flow cytometry (described below).

### 2.4. CD45+ cell isolation from brain

To characterize immune cell populations in the brain during DSS-induced colitis, Miltenyi Biotec’s Adult Brain Dissociation Kit (ABDK, #130-107-677) was used according to manufacturer specifications. Upon harvesting, brains were washed briefly in PBS, cut into roughly 16 small pieces, and put into gentleMACS C-tubes (Miltenyi Biotec, #130-093-237) with dissociation enzymes prepared as instructed. Tissue was subjected to the 37C_ABDK_01 protocol on the gentleMACS Octo-Dissociator with heaters (Miltenyi Biotec, #130-096-427) for dissociation. Brain lysates were filtered through a 70µm filter with Dulbecco’s PBS with calcium, magnesium, glucose, and pyruvate (D-PBS) and pelleted at 300*g* for 10min at 4°C. Lysate was resuspended in D-PBS, Debris Removal Solution was added appropriately, and a layer of D-PBS was deposited gently over the tissue-laden solution. Debris and cell layers were created via centrifugation at 3000*g* for 10min at 4°C with decreased braking, after which the top two phases, containing myelin and cell debris, were discarded. Cells were washed once with D-PBS and re-pelleted at 1000*g* for 10min at 4°C. Red blood cells were removed with Red Blood Cell Removal Solution, diluted 1:10 in double-distilled water, with incubation at 4°C for 10min. Lysis was quenched by adding 10x volume of D-PBS with 0.5% bovine serum albumin (BSA), and cleaned cells were pelleted again at 300*g* for 10min at 4°C.

To enrich for immune cells, the cell suspension was subjected to magnetic separation with CD45+ magnetic bead-labeled antibodies. Cells were resuspended in 90µL of a buffer comprised of autoMACS Rinsing Solution (Miltenyi Biotec, #130-091-022) with 0.5% BSA (BB) and 10µL CD45 MicroBeads (Miltenyi Biotec, #130-052-301) with a 15min incubation at 4°C. Antibody labeling was quenched with the addition of 20x volume BB and cells were pelleted at 300*g* for 10min at 4°C. Labeled cells were resuspended in BB and added to MS columns (Miltenyi Biotec, #130-042-201) in an OctoMACS Separator (Miltenyi Biotec, #130-042-108). Columns were washed three times with BB to elute unlabeled cells, and labeled cells were eluted in BB with the assistance of the plunger. Enriched samples were pelleted at 300*g* for 5min at 4°C, resuspended in PBS, and carried forward to surface epitope staining and flow cytometry (described below).

### 2.5. Flow cytometry

Brain and circulating immune cells were transferred to a V-bottom plate for surface epitope labeling. Samples were centrifuged at 300*g* for 5min at 4°C, washed once in PBS, re-pelleted, and resuspended in an antibody cocktail containing fluorophore-conjugated antibodies (Supp. Table 2) and LIVE/DEAD Fixable Blue stain (ThermoFisher, #L34962) in BD Horizon Brilliant Stain Buffer (BD Biosciences, #563794). Cells were incubated with antibodies for 20min at 4°C in the dark before three washes in FACS buffer containing 0.25mM EDTA, 0.01% NaN_3_, and 0.1% BSA in PBS. Labeled cells were fixed in 1% paraformaldehyde in a phosphate buffer for 30min at 4°C in the dark. Cells were washed three times in FACS buffer and transferred to FACS tubes in FACS buffer for flow cytometry.

For compensation, 1uL of antibody was added to reactive and negative AbC Total Antibody Compensation Beads (ThermoFisher, #A10497) and 0.5uL of LIVE/DEAD Fixable Blue stain was added to ArC Amine Reactive Compensation Beads (ThermoFisher, #A10346). Bead staining occurred at the same time as cell staining and was subjected to the same protocol described above without fixation (e.g., labeled beads remained in FACS buffer while cells were in fixative). Samples were run on a BD FACSymphony A3, capturing either 100,000 single cells or the entire sample. Laser voltages were set with the assistance of SPHERO Supra Rainbow particles (Spherotech, Inc., #SRCP-35-2A) based on settings optimized during the development of this antibody panel. Compensation controls were calculated to ensure successful channel compensation prior to running samples. Raw files were analyzed using FlowJo (BD Biosciences) v.10.9.0 and gates were drawn with the assistance of fluorescence-minus-one controls.

### 2.6. MesoScale Discovery multiplexed immunoassays

MesoScale Discovery (MSD) assays were performed with the recommended protocol from the manufacturer. In this study, the mouse R-Plex CRP assay (#K1525KR-2), mouse U-plex NGAL/LCN2 assay (#K152Z1K-1), and mouse V-Plex Pro-Inflammatory Panel 1 assay (#K15048D0-1) were used. Plasma samples were diluted appropriately for the level of analyte in each assay (1:4 for the Pro-Inflammatory panel, 1:1000 for the CRP assay, 1:100 for the LCN2 assay). The CRP and LCN2 assays require the user to coat the plate with biotinylated capture antibody, which was done by diluting the antibody in Diluent 100 and adding 25µL of the diluted antibody to each well, followed by shaking at 700rpm at room temperature (RT) for 1hr and three washes with PBS with 0.05% Tween-20 (PBS-T). Samples or standard calibrators were added in 50µL duplicates and incubated for 1hr (CRP/LCN2) or 2hr (Pro-Inflammatory). After three washes in PBS-T, SULFO-TAG labeled tracer antibodies were diluted in Diluent 41 and applied at RT for 1hr (CRP/LCN2) or 2hr (Pro-Inflammatory) with shaking at 700rpm. Plates were washed three times with PBS-T, and GOLD Read Buffer B (CRP/LCN2) or a 2X dilution of Read Buffer T (Pro-Inflammatory) was applied just before reading on a MESO QuickPlex SQ 120 workstation.

### 2.7. LBP ELISA

Concentrations of plasma LPS-binding protein (LBP) were examined via Mouse LBP SimpleStep ELISA according to the manufacturer’s protocol (Abcam, #ab269542). Plasma samples were prepared at a 1:16,000 dilution. Standards, antibody cocktail containing detector and capture antibodies, and 1x wash buffer were prepared according to the manufacturer’s protocol. In the plate, 50µL of standard or sample was added to appropriate wells in duplicate. 50µL of antibody cocktail was added to all wells. The plate was then incubated at RT for 1 hour on a plate shaker at 400 rpm. Wells were decanted and washed three times with 150µL 1x wash buffer. 100µL of TMB Development Solution was added to each well and allowed to incubate in the dark for 12 minutes with shaking at 400 rpm. 100µL of Stop Solution was added to each well and placed on the shaker for 1 minute at 400 rpm to mix. Absorbance of 450nm light was recorded with a FLUOstar Omega microplate reader (BMG Labtech). The standard curve was created by plotting the blank-subtracted average LBP standard absorbance against the standard LBP concentrations. Sample LBP concentrations were calculated by interpolating the blank-subtracted average absorbance values against the standard curve, multiplied by the dilution factor.

### 2.8. Protein and nucleic acid extraction

Protein and RNA were extracted from tissue using a TRIzol™-chloroform based method. TRIzol™ (ThermoFisher, #15596018) was added to tissue and lysed using a sterile metal bead in a TissueLyser II (Qiagen). Tissue lysate in TRIzol was then mixed with UltraPureTM phenol:chloroform:isoamyl alcohol (ThermoFisher, #15593049) in a 5:1 ratio. Samples were centrifuged for 15min to form three discrete layers. RNA was taken from the top aqueous layer and passed through a QIAshredder column (Qiagen, #79656). Flow-through was mixed 1:1 with 70% ethanol and passed through a RNeasy Mini column (Qiagen, #74104). After centrifugation, the column was washed once in 700µL RW1 buffer and twice in 500µL RPE buffer according to manufacturer’s instructions. RNA was eluted in nuclease-free water and concentration was measured using a Denovix DS-11 spectrophotometer/fluorometer.

To isolate protein, the DNA layer was removed and discarded as was some of the pink layer until roughly 300µl remained. To this layer, 1mL of methanol was added to precipitate out the protein. After a 10min incubation, samples were centrifuged to pellet the protein. The supernatant was decanted, and pellets were washed in another 500µL of methanol. After centrifugation, the supernatant was again decanted and tubes were air dried to remove residual methanol, about 5 minutes. Protein was resuspended in 1% SDS (w/v) in MilliQ water.

### 2.9. SDS-PAGE & immunoblotting

Protein concentration was measured using a Pierce BCA Protein Assay Kit (ThermoFisher, #23225) according to manufacturer instructions. Within experiments, protein was diluted to an equivalent concentration for each sample in additional 1% SDS solution with 4X Laemmli sample buffer (Bio-Rad, #1610747) and 2-mercaptoethanol. 15µg of protein lysate was loaded into a 4-20% polyacrylamide Criterion TGX stain-free gel (Bio-Rad, #5678095). Electrophoresis was performed using 60V for one hour followed by 125V until finished, about 75 minutes. Samples were transferred to a PVDF membrane (Biorad, #1620175) using a Trans-Blot Turbo Transfer system (Bio-Rad, #1704150) using the preset Mixed MW program.

Signal normalization was achieved using Revert 700 Total Protein Stain (Li-Cor, #926-11011). The stain was imaged on a Li-Cor Odyssey Fc imager with a 2min exposure time. Afterwards, the membrane was cut into sections containing proteins of interest and blocked in 5% dry nonfat milk (w/v) with 0.1% Tween-20 in TBS (TBS-T) for 1hr at room temperature. Primary antibodies (Supplemental Table 2) were applied overnight at 4°C in 5% milk (w/v) in TBS-T. After 3 washes in TBS-T, HRP-conjugated secondary antibodies (Supplemental Table 2) were applied for 1hr at room temperature in 5% milk (w/v) in TBS-T. Membranes were washed 3 times in TBS-T followed by 2 extra washes in TBS to remove excess detergent. To visualize proteins of interest, SuperSignal West Pico (Thermofisher, #34580) or Femto (ThermoFisher, #A34095) were used, depending on the abundance of the targets, according to manufacturer’s instructions. Chemiluminescent reactions were imaged using a Li-Cor Odyssey Fc imager. Immunoblotting images were analyzed using ImageStudio Lite (Li-Cor) and target protein signal intensities were normalized to their respective Total Protein image.

### 2.10. cDNA synthesis & qPCR

Complementary DNA (cDNA) was synthesized from RNA using the ImProm-II Reverse Transcription System (Promega, #A3800) according to manufacturer’s instructions. Briefly, oligo-dT primers were hybridized to at most 1µg of raw RNA in at most 4µL for 5min at 70°C. Then, the reverse transcription master mix, containing 1X ImProm-II reaction buffer, 5mM MgCl2, 0.6mM deoxyribonucleotides, RNase inhibitor, and reverse transcriptase, was added to the RNA with annealed primers. Samples were equilibrated at 25°C for 5min followed by reverse transcription at 42°C for 60min and reverse transcriptase inactivation at 70°C for 15min. Samples were then diluted in nuclease-free water to yield an appropriate working concentration of cDNA to confidently detect gene target of interest, usually 10µg/µL.

Gene expression was quantified using quantitative polymerase chain reaction (qPCR) with SYBR Green-based chemistry. 5µL of cDNA was mixed with 35µL of qPCR master mix, consisting of 1.25µM forward and reverse primers (Supplemental Table 3) obtained from Integrated DNA Technologies (Research Triangle Park, NC, USA), SYBR Green (ThermoFisher, #A46112), and nuclease-free water. This reaction mix was plated into 10µL triplicates. A QuantStudio 5 Real-Time PCR System (ThermoFisher) was used to perform the qPCR. After completion of the assay, technical replicates were pruned based on intra-sample standard deviation (SD) and deviation from the median cycle of threshold (CT) value using a custom R script. If the triplicates showed a SD > 0.3, the replicate with a deviation from the median over 0.5 was discarded. If both the minimum and maximum replicates showed a deviation from the median over 0.5, the sample was discarded. Data were analyzed using the ΔΔCT method, with data being expressed as a fold change from the average ΔΔCT of the reference group.

### 2.11. RNA sequencing – data pre-processing

Prior to RNA sequencing, RNA quality was assessed using a Bioanalyzer 2100 (Agilent) with the RNA 6000 Nano Kit (Agilent, #5067-1511) according to manufacturer’s instructions. Samples with RIN < 7, 260/230 < 1.0, and 260/280 < 1.8 were excluded. For each sample, 1µg was diluted to 20ng/µL in nuclease-free water and shipped to LC Sciences for cDNA library preparation and sequencing. Their workflow is described below.

Messenger RNA was enriched from total RNA using DynabeadsTM Oligo(dT)25 (ThermoFisher, #61005) with two rounds of purification. The mRNA was fragmented using divalent cations under 94°C for 5-7min using the NEBNextTM Magnesium RNA Fragmentation Module (New England Biolabs, #E6150S). RNA fragments were reverse-transcribed using SuperScriptTM II Reverse Transcriptase (Invitrogen, #1896649) into cDNA. The cDNA was used to synthesize U-labeled second-stranded DNAs with E. coli DNA polymerase I (New England Biolabs, #M0209), RNase H (New England Biolabs, #M0297), and dUTP Solution (ThermoFisher, #R0133). An A-base was added to the blunt ends of each strand. DNA was ligated to dual-index adapters, containing a T-base overhang for ligation on the A-tailed DNA fragments. Size selection was performed with AMPureXP beads (Beckman Coulter, #A63882). DNAs were treated with heat-labile UDG enzyme (New England Biolabs, #M0280), and ligated products were amplified with PCR with initial denaturing at 95 for 3min, 8 cycles of denaturation at 98 for 15sec, annealing at 60 for 15sec, and extension at 72 for 30s, followed by a final extension at 72 for 5min. The average insert size for cDNA libraries was between 250-350bp. Finally, 2×150bp paired-end sequencing was performed on an Illumina NovaseqTM 6000 according to the manufacturer-recommended protocol.

Reads were filtered to yield only high-quality reads with Cutadapt, using the following parameters: 1) reads containing adapters were removed; 2) reads containing polyA and polyG were removed; 3) reads containing more than 5% of unknown nucleotides; 4) reads with more than 20% low-quality bases (Q-value < 20) were removed. Sequence quality was verified using FastQC, considering the Q20, Q30, and GC-content of the cleaned data. Reads were mapped to a reference mouse genome (GRCmm39, from GENCODE) using STAR (Dobin et al., 2013) and raw count matrices were generated using ‘featureCounts’ from the Subread package (Liao et al., 2013) using the genome’s accompanying annotation set.

### 2.12. RNA sequencing – quality control

Brain and colon datasets were treated separately for all subsequent analyses. In each dataset, genes were discarded if fewer than eight samples had more than 0.5 counts per million (CPM) with the edgeR package. Variance stabilized data were carried forward to co-expression analysis. Inter-sample distances were calculated and outlier samples were removed based on visual inspection of the sample dendrogram within each of the six data sets, as recommended by the WGCNA tutorial (https://horvath.genetics.ucla.edu/html/CoexpressionNetwork/Rpackages/WGCNA/Tutorials/). The WGCNA package (Langfelder and Horvath, 2008) includes feature pruning based on a lack of variance and low number of counts, which was used to further discard low quality genes from the data sets. After gene and sample removal, datasets were rerun through DESeq2 (Love et al., 2014) to generate re-normalized and variance stabilized counts. Gene quality was re-checked with the steps described above. If more genes were flagged for removal, the dataset would be re-normalized again until all genes passed this quality check. The data generated for this study are accessible through NCBI’s Gene Expression Omnibus via the GEO Series accession number GSE239820 (https://www.ncbi.nlm.nih.gov/geo/query/acc.cgi?acc=GSE239820).

### 2.13. RNA sequencing – differential expression analysis

After ensuring the retention of only high-quality genes and samples, the brain dataset was split into cortex, hippocampus, midbrain, and striatum subsets. Differential expression was conducted in each regional subset using DESeq2, including treatment group as a fixed effect, with default settings. Differentially expressed genes, defined as having an absolute log2-transformed fold-change > 1 and Benjamini-Hochberg adjusted *p*-value < 0.05, were generated at each DSS group relative to controls for each brain region, resulting in 24 sets of differentially expressed genes. Volcano plots were created to visualize these sets, and Venn diagrams were created at each treatment group to compare the overlap of differential expression between brain tissues.

### 2.14. RNA sequencing – gene set proximity analysis (GSPA)

To further compare the lists of differentially expressed genes due to DSS in different brain regions, a GSPA was performed (Cousins et al., 2023). This analysis extends the classical gene set enrichment analysis framework developed by the Broad Institute (Subramanian et al., 2005) by assigning protein-coding genes to an embeddings space built from high-confidence protein-protein interactions from the STRING repository (Szklarczyk et al., 2017). Enrichment scores, and the null distributions from which they are derived, integrate the interconnectedness of genes, providing more specific significance estimates. To perform this analysis with our data, DESeq2 results from each group in each tissue were generated and the effect size was shrunk based on data quality using the “apeglm” shrinkage estimator (Zhu et al., 2019) for more accurate ranking. Genes were converted to human symbols via the interconversion of ENTREZ identifiers and were discarded if adjusted *p*-values were not calculated due to the presence of extreme outliers based on Cook’s distance, if a human homologue was not found, or if the gene was not found in the GSPA’s high-confidence STRING-derived embeddings list. This analysis was performed according to the tutorial at https://github.com/henrycousins/gspa using the human MSigDB Hallmark list of gene sets (Liberzon et al., 2015), and normalized enrichment scores are shown. The FDR-corrected *p*-values were computed based on a single gene list, so these values were further Bonferroni-adjusted to account for the 24 comparisons being made here.

### 2.15. RNA sequencing – weighted gene correlation network analysis (WGCNA)

To evaluate global transcriptomic perturbations in the brain as a result of a leaky gut, adjacency networks using Pearson correlations were created within each of the four brain data sets using a soft thresholding power of 6, chosen with assistance from the WGCNA’s package ‘pickSoftThreshold’ function. Signed hybrid networks were employed in this study, where positive correlations were raised to the soft thresholding power and negative correlations were set to zero. A topological overlap matrix (TOM) was calculated from each adjacency network. The midbrain, striatum, and hippocampus TOMs were scaled on the 95^th^ quantile of the cortex TOM as the cortex was arbitrarily selected as the “first” dataset of the four. The minimum topological overlap weight for each node was taken as the consensus TOM between all data sets. The consensus TOM was then converted to a distance matrix by subtracting the TOM from 1, and this distance measure was used to identify gene co-expression modules using an average linkage hierarchical clustering strategy. Modules were merged if they were more than 75% correlated. Gene module membership (kME) was determined by correlating the gene counts with the eigengene. To compute the meta-kME for a gene across all databases, these correlation coefficients were transformed to Fisher’s Z-values, which were summed and divided by the square root of the number of data sets. Global transcriptomic perturbations in the colon were evaluated similarly using the distal and proximal colon data sets. Soft thresholding power was set to 7, and the proximal colon TOM was scaled on the 95th quantile of the distal colon TOM. Gene level information including kME and meta-kME can be found in Supplementary File 1 (colon) and Supplementary File 2 (brain).

After consensus TOM construction and consensus module identification, the individual datasets were analyzed separately with the same soft power threshold, network type, and clustering strategy to identify tissue-specific gene modules. To assess how well the tissue-specific co-expression modules were preserved in the respective consensus network, Fisher’s exact test was performed between each possible pair of specific and consensus modules. A Bonferroni correction was manually applied to this group of tests. Results from this analysis are presented in a large tile matrix, where each tile displays the number of overlapped genes and are colored by whether the adjusted *p*-value from the contingency table analysis is below 0.05.

To identify interesting gene modules in colon, module function was inferred through gene set enrichment analyses (described below). In brain, eigengenes were correlated with measures of disease severity, circulating proteins, and key colon consensus module eigengenes. To account for a large set of correlations occurring simultaneously, *p*-values from these relationships were adjusted with Bonferroni’s correction. The WGCNA package also computes correlations between each gene and each external trait within each tissue, designated gene-significance (GS), and the meta-GS by summing the Fisher transforms of the correlation coefficients divided by the square root of the number of datasets. Gene level information including GS, meta-GS, and the respective *p*-values can be found in Supplementary File 1 (colon) and Supplementary File 2 (brain).

### 2.16. RNA sequencing – gene set enrichment of WGCNA-identified co-expression modules

Functional annotation of gene co-expression modules was performed using the anRichment R package (https://horvath.genetics.ucla.edu/html/CoexpressionNetwork/GeneAnnotation/). This package interfaces fluently with the output of the WGCNA package and allows for rapid calculation of the enrichment of supplied reference gene sets. A comprehensive collection of reference gene sets was built by combining all the gene lists included in the Gene Ontology (GO) knowledgebase’s Biological Process category (Ashburner et al., 2000; Gene Ontology Consortium et al., 2023), Kyoto Encyclopedia of Genes and Genomes (KEGG) (Kanehisa et al., 2010), BioCarta database (Rouillard et al., 2016), WikiPathways database (Martens et al., 2021; Pico et al., 2008), REACTOME database (Gillespie et al., 2022), and the Hallmark gene sets from the mouse Molecular Signatures Database (MSigDB) (Liberzon et al., 2015, 2011). Enrichment scores, gene overlap, and corrected enrichment *p*-values from this large gene set collection were calculated for each module within a consensus network. This analysis was performed with ‘background = “given”’ to use every gene found in our dataset as the background rather than extra genes not discovered here that may be present in the reference gene sets.

### 2.17. RNA sequencing – module preservation analysis with publicly available datasets

To assess the reproducibility of molecular features of colitis, we performed a module preservation analysis using the WGCNA package in R (Langfelder et al., 2011). This analysis evaluates whether the connectivity profile of a module found in a reference network, which in this study was either our distal or proximal colon network, is similar to that of a test network if genes are assigned to the same modules. To determine if preservation is better than should be expected due to chance, module assignment in the test network is randomly permuted 200 times and network statistics are estimated which are compared to the observed statistics arising from the module assignment of interest, yielding *Z*-score transformations of key statistics and *p*-value estimations.

From this analysis, we show *Zsummary*, a composite statistic that summarizes several individual *Z*-statistics. This summary score takes the average of density-based preservation, interpreted as the degree to which module nodes remain highly connected, and a measure of connectivity-based preservation, interpreted as the degree to which patterns of connectivity are similar between networks. According to simulation studies, there is little support for module preservation when *Zsummary* < 2, weak-to-moderate support for preservation when 2 < *Zsummary* < 10, and strong support for preservation when *Zsummary* > 10.

We used three datasets to evaluate the reproducibility of gene programs discovered here. The first is GSE131032, a similar time-series study where female C57Bl6/J mice were given 2.5% DSS for 7d followed by 7d of recovery, and mid-colon tissue was collected every other day (Czarnewski et al., 2019). The second is GSE168053, a time-series study where C57Bl6/J mice received 3% DSS for 6d followed by 14d of recovery, and several sections of colon were harvested at 0d, 3d, 6d, 9d, 14d, or 20d (Liu et al., 2022). This study included separate samples for distal and proximal colon, permitting us to evaluate not only the preservation of modules but examine the robustness of region-specific processes during DSS-induced colitis. The third dataset was GSE210405, a small study where mice received 3% DSS for 6d and mid-colon tissues were harvested every other day (unpublished; Zheng et al., 2022). Normalized counts were used as provided with no additional quality control or normalization measures on our side. All code needed to reproduce the transcriptomic analyses here can be found at https://github.com/jakesboles/Boles_et-al_DSS_time-series.

### 2.18. Statistical analysis

Statistical analysis was performed in R v.4.2. Assays involving a single variable in Cohort 1, including measuring circulating immune markers and qPCRs, were analyzed using a one-way ANOVA using the ‘aov_ez’ function from the Afex package (https://cran.r-project.org/web/packages/afex/index.html). The Performance package was used to check homogeneity of variance between groups. When this assumption was met (i.e., Levene’s test was not significant), *post hoc* comparisons were made with Tukey’s correction for multiple comparisons using the Emmeans package (https://cran.r-project.org/web/packages/emmeans/index.html). When the assumption of homogeneity of variance was violated, a new model was fitted using ‘anovaOneW’ from the JMV package (https://cran.r-project.org/web/packages/jmv/index.html). This function was called with ‘welchs = TRUÈ to use Welch’s correction to the degrees of freedom when inter-group variances are heterogeneous and ‘phMethod = “gamesHowell”’ to use the Games-Howell correction for multiple comparisons during *post hoc* testing. Assays involving two independent variables in Cohort 1, including immunoblotting analysis of tight junctions in different colon segments, were analyzed using a two-way ANOVA using the Afex package. As colon segment represented a within-subjects factor, sphericity was assessed using the Performance package and the Geisser-Greenhouse correction was applied as needed.

Analysis of RNA sequencing count data, which included both tissue and treatment group was independent factors, was done using a set of mixed-effects models using the “lmer” function from the lme4 package (https://cran.r-project.org/web/packages/lme4/index.html) to handle the random loss of samples due to RNA or sequencing quality, unrelated to treatment group or brain region. A “null” model was fit that included the fixed effect of brain region and the random effect of mouse. A “main effects” model was fit that included the fixed effects of brain region and treatment group and the random effect of mouse. To evaluate whether treatment group significantly affected the variance in the sample, these two models were compared with base R’s “anova” function. A third “interaction” model was fit that included both main effects and the random effect described in the main effects model and the interaction between brain region and group. Base R’s “anova” was called with this interaction model and the main effects model to evaluate whether the interaction between variables was significant.

To succinctly visualize significant differences between groups, compact letter displays (CLDs) were generated for analyses that revealed a significant main effect of treatment group using the Biostat package’s ‘make_cld’ after running ‘posthoc_anovà (https://gegznav.github.io/biostat/) on the model when homogeneity of variance was violated in one-way designs or Multcomp’s ‘cld’ (https://cran.r-project.org/web/packages/multcomp/index.html) when homogeneity of variance was met or in multi-factor designs. In figures containing multiple tissues, the CLD was created within each tissue, meaning groups were not compared across separate brain regions or colon segments. In all figures, groups that do not share a letter are statistically significantly different (adjusted *p* < 0.05). Conversely, groups that do share a letter are not significantly different (adjusted *p* > 0.05).

In Cohort 2, most cell population statistics were analyzed with a *t*-test or non-parametric alternative. JMV’s ‘ttestIS’ was used, which allows the simultaneous assessment of *t*-test assumptions and hypothesis testing. This function was called with “norm = T” to perform the Shapiro-Wilk test of normality and “eqv = T” to perform Levene’s test for homogeneity of inter-group variance. When both tests showed *p* > 0.05, Student’s *t* is reported. When the Shapiro-Wilk test was significant (*p* < 0.05), the non-parametric Mann-Whitney *U* is reported. When Levene’s test was significant (*p* < 0.05), Welch’s *t* is reported. When both tests were significant, Welch’s *t* was reported as the Mann-Whitney alternative assumes the distributions of each group are equivalent. In certain cases where the relative abundance of cell subtypes were deemed dependent on each other, the data were treated with a two-way ANOVA, where cell subtype abundance was the within-subjects factor and DSS was the between-subjects factor, and the Geisser-Greenhouse correction was used. Here, pairwise comparisons were made at each cell type separately with the Emmeans package and *p*-values were adjusted with Bonferroni’s correction. All visualizations were created in R with base R plotting language or the ggplot2 package (https://cran.r-project.org/web/packages/ggplot2/index.html).

## 3. Results

### 3.1. The severity of DSS-induced colitis peaks immediately after DSS withdrawal

To characterize the brain’s response to a “leaky” and inflamed gut and the associations between disturbances along the gut-brain axis during intestinal inflammation, mice received 2.5% DSS (w/v) in their drinking water according to varied schedules, yielding seven total groups: untreated controls, 5d DSS, 7d DSS, 7d DSS + 2d recovery, 7d DSS + 5d recovery, 7d DSS + 7d recovery, and 7d DSS + 14d recovery (Fig. 1A). This time-series design allowed us to define temporally distinct molecular signatures in the brain and understand how these are related to discrete phases of damage and inflammation in the intestine. Body weight loss occurred over the 7d of DSS treatment and continued for 2d after which mice steadily regained weight to baseline status (Fig. 1B). A daily disease index, combining weight loss, stool consistency scores, and the presence of rectal bleeding/fecal occult blood, showed worsening and recovery along the same timeline (Fig. 1C). Upon euthanasia, the spleen and colon were measured and weighed as coarse indexes of disease severity and systemic inflammation. DSS treatment induced significant changes in spleen weight (Fig. 1D) and colon length (Fig. 1E). Spleens were enlarged after 7d until after 7d of recovery (Fig. 1D), indicative of a systemic immune response, and colons were shorter after 7d of DSS, reflective of intestinal injury (Fig. 1E). To confirm the efficacy of DSS administration in denuding the intestinal epithelium and establish phases of damage and repair, we examined intestinal tight junction protein content via western blot. The loss of claudin-2 was rapid and persistent, showing a similar pattern of loss in both colon segments (Fig. 1F-G). The loss of occludin was also rapid and persistent, although the distal colon was more persistently affected by DSS (Fig. 1H-I). Loss of ZO-1 was transient, worse in the distal colon, and demonstrated a recovery after a week of untainted drinking water with a reinduction of loss after two weeks of untainted drinking water (Fig. 1J-K). Together, these data demonstrate a successful induction of colitis with DSS, characterized by a typical exacerbation of disease indexes and loss of intestinal tight junction proteins.

### 3.2. DSS administration induces temporally-defined intestinal immune activation

To define intestinal molecular processes at play during colitis and determine which processes are reproducible, we performed a consensus co-expression analysis in distal and proximal segments of the colon harvested here using the WGCNA R package (Langfelder and Horvath, 2008). A signed hybrid adjacency network was chosen, and the beta power for the network was set to 7 on the recommendation of selecting the lowest power to achieve a model fit with R^2^ > 0.8 (Fig. S1A-D). The consensus topological overlap was calculated as described (Langfelder and Horvath, 2008), and genes were clustered into co-expression modules which were merged if they were 75% correlated, leaving 58 unique modules (Figure S1E). Overall, the global profile of module assignment was well preserved between distal and proximal segments (Fig. S2A-D). Furthermore, set-specific modules identified by building separate co-expression networks for each segment showed significant overlap with consensus modules (Fig. S2E-F), suggesting that co-expression patterns are preserved from one part of the colon to the other. To identify disease-associated modules, a gene set enrichment analysis was conducted using the anRichment R package (https://horvath.genetics.ucla.edu/html/CoexpressionNetwork/GeneAnnotation/), employing gene sets from the GO (Ashburner et al., 2000; Gene Ontology Consortium et al., 2023), KEGG (Kanehisa et al., 2010), MSigDB (Liberzon et al., 2015; Subramanian et al., 2005), BioCarta (Rouillard et al., 2016), REACTOME (Gillespie et al., 2022), and WikiPathways (Martens et al., 2021; Pico et al., 2008) databases.

Given the prevailing hypothesis that inflammation in the intestine is the driver of gut-to-brain propagation of inflammation and/or injury, we first examined the expression profiles of immunological gene signatures. Moreover, we evaluated whether these gene signatures were preserved in three other datasets (GSE131032, from KTH Solna, Stockholm, Sweden (Czarnewski et al., 2019); GSE168053, from University of Chicago, Chicago, IL (Liu et al., 2022); GSE210405, from Zheng et al., 2022, Shandong University, Jinan, China) procured from intestinal RNA-sequencing during a DSS dosing scheme to determine whether these signatures are reproducible or depend on experimental context. The ‘blueviolet’ consensus module showed strong enrichment of pathways associated with neutrophil activation and innate immunity (Fig. 2A). In our dataset, this module peaked alongside disease severity and remained up-regulated after the withdrawal of DSS, while GSE131032 and GSE168053 demonstrated a reversible activation and GSE210405 showed a corroborating rise during DSS administration (Fig. 2B). Genes such as *S100a9*, *Cxcl2*, *Mmp9*, and *Lcn2* were strong members of this module, reflecting the enrichment of neutrophil systems (Fig. 2C). This module showed at least moderate support for preservation in all three datasets, suggesting that this module is authentic during DSS-induced colitis (Fig. S3). The ‘darkseagreen3’ module displayed an enrichment of myeloid cell programs, including phagocytosis (Fig. 2D), and showed a similar expression profile as the ‘blueviolet’ neutrophil module: our dataset suggested activation during DSS with perseverance post-withdrawal, while GSE131032 and GSE16805 demonstrated transient peaks and GSE210405 showed the expected increase during DSS (Fig. 2E). This module included genes such as *Tyrobp*, *Tgfb1*, *Apoe*, and *C1qa* (Fig. 2F), and showed weak-to-moderate support for preservation in GSE131032 and GSE168053 and strong support in GSE210405 (Fig. S3). The ‘darkorange’ module was enriched with genes associated with cell chemotaxis and the immune system (Fig. 2G) and its expression mirrored that of the two modules described above (Fig. 2H). Hub genes for this module included several genes associated with the crosstalk between tissue barriers and immune cells, including *Pecam1*, *Selp*, and *Ccr1* (Fig. 2I), and this module showed strong support for preservation in all datasets (Fig. S3), suggesting that the mobilization of immune cells in the gut during colitis is reproducible. According to these three modules, myeloid cell reactivity and migration appear to be reproducible phenomena during DSS-induced colitis, and our dataset showed a uniquely persistent activation of these gene signatures during and after treatment.

**Figure 2:**
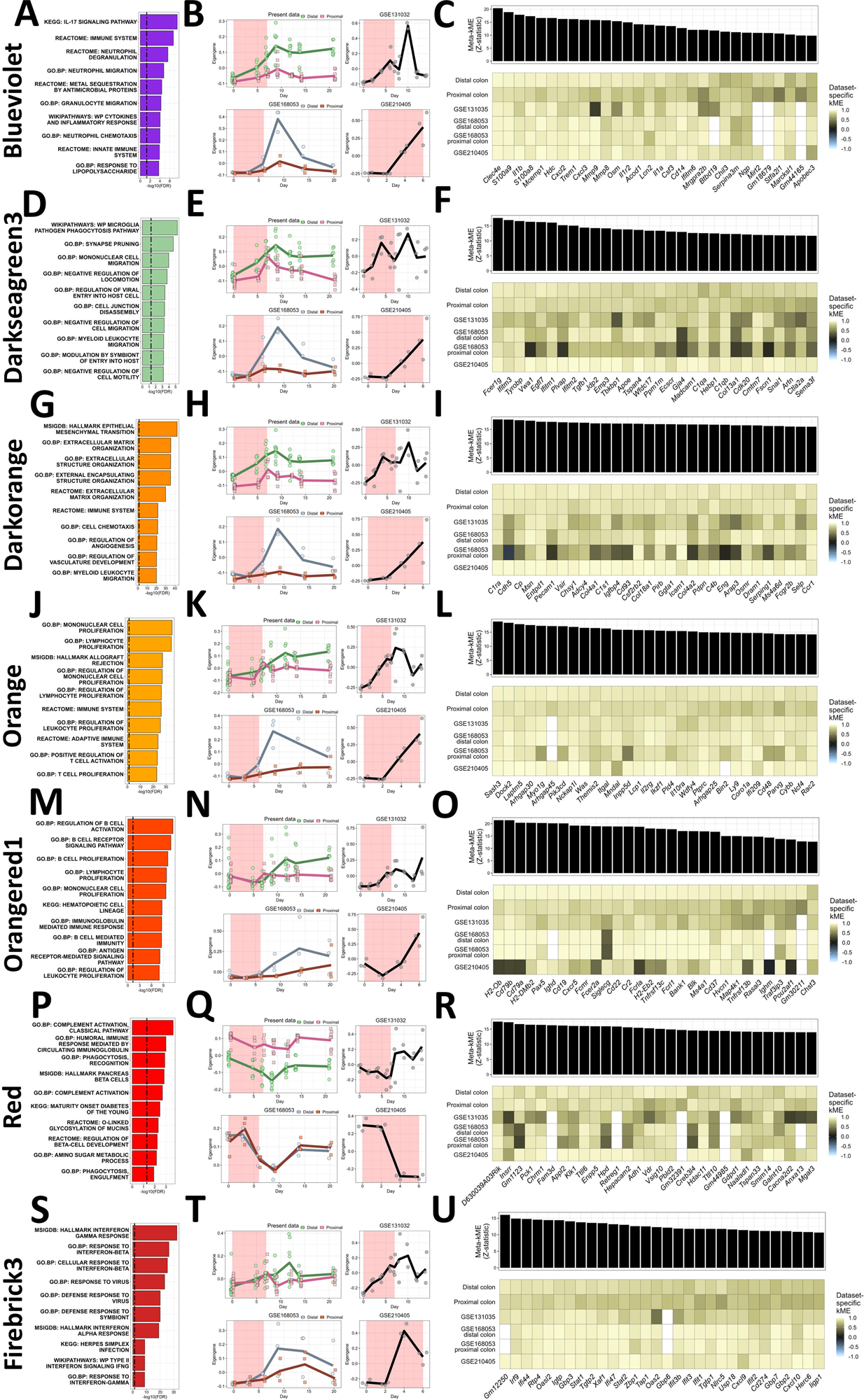
Reproducible and study-specific intestinal immune gene programs. Gene set enrichment results identifying immunological gene modules are shown in **A, E, I, M, Q, U,** and **Y.** The top 10 gene sets are displayed, ranked on their unadjusted *p-* value. Bar heights represent the -log10 transform of the FDR, and the dashed line reflects FDR = 0.05. Module eigengenes for our colon datasets, GSE131032, distal and proximal datasets from GSE168053, and GSE210405 are shown in **B, F, J, N, R, V,** and **Z.** The x-axis represents the time during each respective study, and the red shading signifies when DSS was administered. The line connecting groups represents the mean eigengene in that group. The top 30 module members are shown in **C, G, K, O, S, W,** and **AA.** A meta-module membership statistic is shown as a bar for each gene above the corresponding dataset-specific correlation with the eigengene shown in a heatmap.

The ‘orange’ module was enriched with mononuclear cell proliferation pathways and adaptive immune activation pathways (Fig. 2J) with a dataset-specific expression profile similar to those presented already (Fig. 2K). Hub genes in this module included *Sash3*, *Itgal*, *Lcp1*, *Il2rg*, *Ptprc*, and *Cd48* (Fig. 2L). This module was also strongly preserved in all datasets (Fig. S3), suggesting that the involvement of adaptive immunity during DSS-induced colitis is authentic. The ‘orangered1’ module was enriched in B-cell functions, such as immunoglobulin signaling and antigen presentation (Fig. 2M). This module showed a less drastic and more variable activation in all datasets, including ours, although each dataset showed an up-regulation during or soon after colitis (Fig. 2N). This module was marked by many genes associated with B-cells broadly or antigen presentation, such as *H2-Ob*, *Cd19*, both CD79 antigen genes, and *Ighm* (Fig. 2O). This module was strongly preserved in GSE168053 and only moderately/weakly preserved in GSE131032 and GSE210405 (Fig. S3), supporting a context-specific activation of B-cells in the DSS model. The ‘red’ module was enriched with gene sets attributed to complement signaling, immunoglobulin signaling, and phagocytosis (Fig. 2P) but was downregulated during colitis in our dataset, GSE168053, and GSE210405 (Fig. 2Q), seemingly in contrast to the ‘orangered1’ and ‘orange’ modules. Upon further inspection, *Mgat3*, a transcriptional activator of IgG molecules (Klasić et al., 2018), and genes encoding for IgG1 and many IGHV molecules were found in this module (Supplementary Files 1 and 3), which corroborates an earlier study demonstrating a specific loss of IgG in the intestine during DSS (Stevceva et al., 2001). This module may also reflect intestinal health in general, with the intestine-specific annexin *Anxa13* and nutritional response elements such as *Pck1* and *Fam3d* as strong module members (Fig. 2R). This module was strongly preserved in GSE168053 and GSE210405 and weakly/moderately preserved in GSE131032 (Fig. S3). Finally, our analysis highlights the ‘firebrick3’ module, enriched with interferon signaling pathways, which was activated in all three public datasets while only weakly and variably activated in ours (Fig. 2S-T). Hub genes in this module consisted of many interferon-responsive elements like *Irf9*, *Ifi47*, *Ifit3*, and *Ifit1,* as well as chemokines *Cxcl9* and *Cxcl10* (Fig. 2U). Notably, this module was strongly preserved in GSE168053 and GSE210405 and weakly/moderately preserved in GSE131032 (Fig. S3), suggesting that this co-expression module is robust in DSS-induced colitis but was not evoked in our experimental context as in others. In summary, we detected a complex immune activation in the colon of the DSS model, marked by the recruitment of myeloid and adaptive immune cells, a polarization of antibody repertoire, and interferon signaling, all of which may depend on the immune context and setting in which mice are dosed.

### 3.3. DSS induces a stress response and metabolic dysfunction in the colon, followed by the induction of robust repair mechanisms

We examined the nature and reproducibility of cellular stress, metabolic dysfunction, and tissue repair in the colon after colitis induction, and expected that these processes may display sensitivity to technical differences between experiments. We identified four co-expression modules in the gut that reflect cellular stress, shown by their enrichment for gene sets associated with apoptosis, mitochondrial membrane permeabilization, and misfolded protein responses (Fig. S4A-D). All four of these modules were up-regulated concomitant with peak colitis in our dataset and were up-regulated in the three other datasets examined here during/after colitis (Fig. S4E-H). Yet, the connectivity of each of these modules is preserved to different degrees in each dataset. GSE210405 demonstrated strong preservation of 3/4 stress modules, GSE131032 demonstrated strong preservation of 2/4 modules, and GSE168053 showed only weak-to-moderate support for all (Fig. S3). Strong members of the ‘coral2’ and ‘coral3’ modules show similar membership in other datasets, while several hub genes for the ‘white’ and ‘thistle2’ modules, enriched for protein folding pathways, were likely poor module members in these other datasets, reflected by their low or negative kME (Fig. S4I-L). Cellular stress appears to be a unifying feature of DSS-induced colitis due to the unified up-regulation of these systems in several independent studies. Still, the specific molecules involved in protein processing during colitis appear to differ between experiments.

Our consensus network approach revealed seven modules enriched for gene sets associated broadly with metabolic processes, including amino acid synthesis and the citric acid cycle (Fig. S5A-G). All but the ‘darkslateblue’ module were downregulated in our dataset during disease, while both GSE168053 and GSE210405 displayed downregulation of all seven modules during DSS-induced colitis. GSE131032 indicated no clear regulation during DSS-induced colitis followed by an up-regulation after DSS is withdrawn, suggesting that intestinal energetic balance may be highly influenced by experimental design or setting (Fig. S5H-N). Module connectivity also appeared to differ between experiments – only the ‘brown’ module, associated with lipid oxidation and catabolism, was strongly preserved in all three datasets, and its members showed strong membership in all datasets (Fig. S3, S5O). The ‘coral,’ ‘darkolivegreen4,’ ‘lightpink3,’ and ‘coral1’ modules showed weak/moderate support for preservation in all datasets and poor preservation of module members, the ‘indianred4’ module is strongly preserved only in GSE168053, and the ‘darkslateblue’ module was strongly preserved in GSE131032 and GSE210405 (Fig. S3, S5P-U). The degree of preservation is overall corroborated by module membership across datasets. Thus, although the broad regulation of metabolic systems is present during colitis in all datasets examined here, the direction of this regulation and the molecular systems recruited differ between experiments.

Finally, our analysis revealed four co-expression modules associated with cell division, DNA replication, and protein synthesis, likely involved in the repair of the intestine after the induction of a leaky gut (Fig. S6A-D). Accordingly, these processes were up-regulated after the withdrawal of DSS in our dataset, GSE131032 and GSE168053 (Fig. S6E-H). GSE210405 did not include a recovery phase, but our analysis suggests an activation of these systems near the end of colitis in this experiment (Fig. S6E-H). All four of these modules were strongly preserved in all datasets (Fig. S3), and the top module members for each showed strong membership in all datasets as well (Fig. S6I-L). Although the induction of an immune response, the nature of the stress response, and metabolic dysfunction may differ between studies using the DSS model, the engagement of mitosis, ribosome formation, and gene expression in the colon appear to mark post-DSS recovery authentically.

### 3.4. A systemic immune response emerges concurrently with peak colitis

Current evidence suggests that inflammatory signals may propagate from the gut to the brain via the circulation (Houser and Tansey, 2017; Jackson et al., 2022), so plasma was collected from mice at all time points described in Fig. 1A, and circulating concentrations of various cytokines and inflammatory markers were measured. DSS treatment significantly affected circulating LBP (Fig. 3A), CRP (Fig. 3B), and LCN2 (Fig. 3C), with each showing significant elevation after 2d of untainted water before returning to baseline levels. DSS affected circulating TNF, which peaked at the same time point, although TNF concentrations remained statistically not different from peak or baseline levels throughout the rest of the schedule (Fig. 3D). Circulating KC/GRO rose above baseline levels at 7d of DSS and peaked 2d after withdrawal but remained elevated 5d after withdrawal (Fig. 3E). Circulating IL-6 showed a significant association with DSS treatment but Games-Howell *post hoc* tests revealed no significant pairwise differences between groups (Fig. 3F). Finally, DSS treatment significantly affected neither IL-10 (Fig. 3G) nor IL-5 levels in circulation (Fig. 3H).

**Figure 3:**
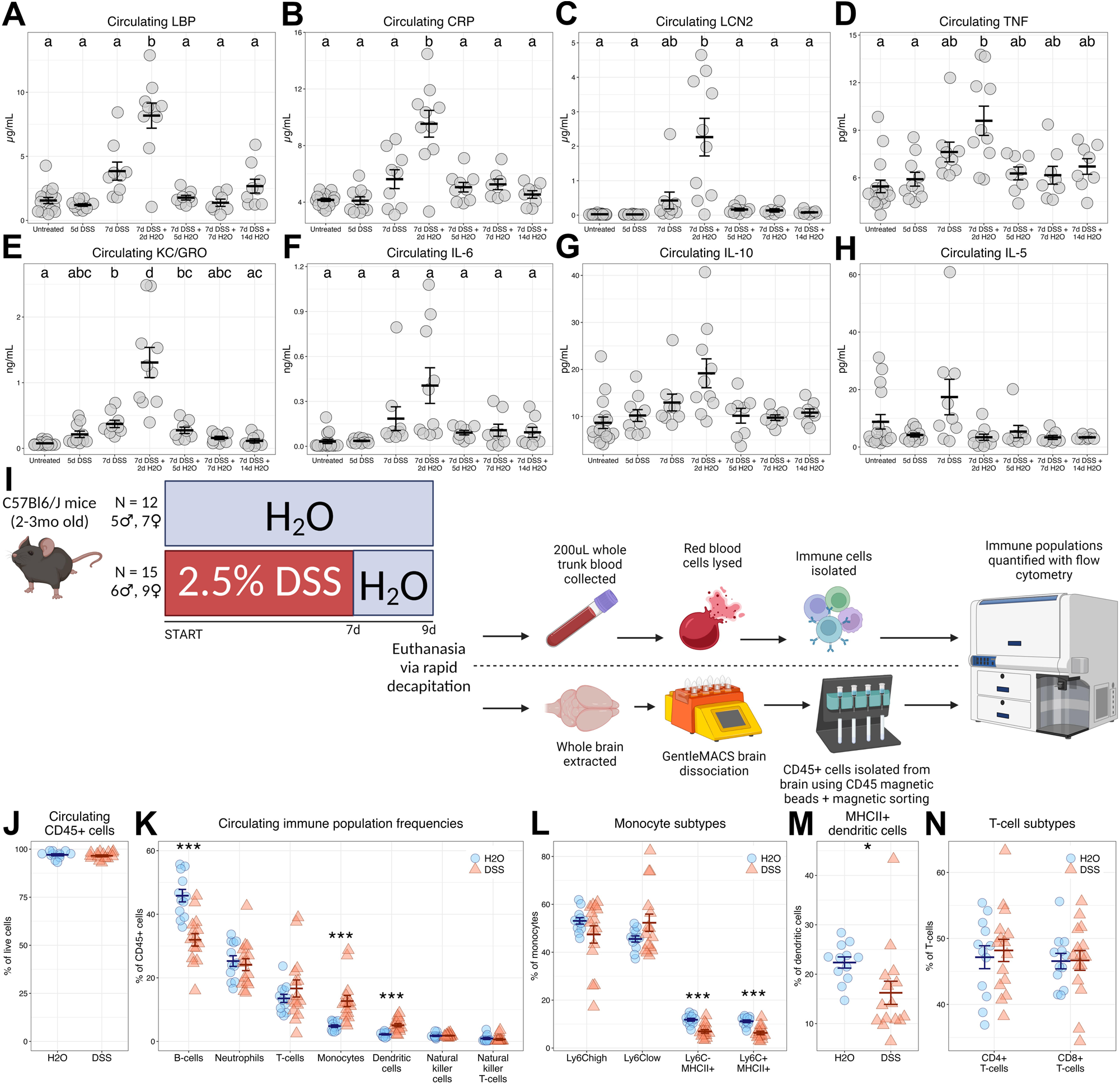
Systemic inflammation peaks concomitant with maximal colitis severity and is marked by innate immune activation. Circulating LBP (**A**; one-way ANOVA: Welch’s *F*[6, 24.298] = 10.651, *p* < 0.001), CRP (**B**; one-way ANOVA: Welch’s *F*[6, 22.780] = 7.069, *p* < 0.001), LCN2 (**C**; one-way ANOVA: Welch’s *F*[6, 22.584] = 6.812, *p* < 0.001), TNF (**D**; one-way ANOVA: Welch’s *F*[6, 25.513] = 3.435, *p* = 0.013), KC/GRO (**E**; one-way ANOVA: Welch’s *F*[6, 21.902] = 12.111, *p* < 0.001), IL-6 (**F**; one-way ANOVA: Welch’s *F*[6, 23.108] = 4.427, *p* = 0.004), IL-10 (**G**; one-way ANOVA: Welch’s *F*[6, 25.835] = 2.018, *p* = 0.100), and IL-5 (**H**; one-way ANOVA: Welch’s *F*[6, 24.674] = 1.696, *p* = 0.164) were measured at all time points described in Figure 1. In these figures, groups that share a letter were not statistically significantly different (*p* > 0.05) according to *post hoc* testing. **I** Timeline and methods of follow-up experiment performed to characterize circulating and brain immune cell populations during peak disease with DSS. **J** Counts of CD45+ cells in circulation after DSS (Student’s *t*[25] = - 0.712, *p* = 0.483). **K** Proportions of specific immune populations (two-way ANOVA: main effect of DSS – *F*[1, 25] = 1.768, *p* = 0.196; main effect of population – *F*[2.673, 66.827] = 162.350, *p* < 0.001; interaction – *F*[2.673, 66.827] = 9.331, *p* < 0.001). *Post hoc* testing was conducted within each cell type and *p*-values were adjusted with Bonferroni’s correction. **L** Monocyte subtypes, discriminated by Ly6C and MHC-II surface staining (two-way ANOVA: main effect of DSS – *F*[1, 25] = 20.453, *p* < 0.001; main effect of population – *F*[1.034, 25.848] = 179.009, *p* < 0.001; interaction – *F*[1.034, 25.848] = 2.908, *p* = 0.099). *Post hoc* testing was conducted within each cell type and *p*-values were adjusted with Bonferroni’s correction. **M** Percentage of circulating dendritic cells that are MHC-II+ (Mann-Whitney U = 31.5, *p* = 0.005). **N** T-cell subtypes, discriminated by CD4 and CD8 surface staining (two-way ANOVA: main effect of DSS – *F*[1, 25] = 0.099, *p* = 0.756; main effect of subtype – *F*[1, 25] = 362.027, *p* < 0.001; interaction – *F*[1, 25] = 0.099, *p* = 0.756). All summary statistics shown are mean ±LSEM. *p < 0.05, ***p < 0.001.

In a follow-up experiment, a separate cohort of mice was dosed according to Fig. 3I. This experiment was designed to elicit peak colitis, informed by disease indexes in Fig. 1B-E and circulating immune markers in Fig. 3A-E, to enable us to examine alterations in circulating immune cell populations that may drive the systemic immune response detected here. The gating strategy used to identify cell populations with flow cytometry is illustrated in Fig. SF5. At peak colitis, there were no differences in overall CD45+ cell abundance in circulation (Fig. 3J, Fig. S7B). Of these CD45+ cells, peak colitis was associated with a lowered abundance of B-cells (Fig. 3K, Fig. S7C) while monocytes (Fig. 3K, Fig. S7D) and dendritic cells (Fig. 3K, Fig. S7E) both made up larger proportions. The abundances of neutrophils (Fig. 3K, Fig. S7F), T-cells (Fig. 3K, Fig. S7G), natural killer cells (Fig. 3K, Fig. S7H), and natural killer T-cells (Fig. 3K) in circulation were not affected by colitis, although we report a small but significant decrease in the counts of natural killer T-cells in circulation with DSS (Fig. S7I). Of the total monocyte population, DSS elicited no differences in Ly6C+ cell frequency, while both populations of MHC-II+ monocytes were present with lower frequency (Fig. 3L). Meanwhile, both Ly6C+ monocyte subtypes showed higher counts with DSS (Fig. S7J-K) while counts of both MHC-II+ monocyte subtypes were unaltered by DSS (Fig. S7L-M). Similarly, fewer dendritic cells were MHC-II+ during colitis (Fig. 3M), while raw counts of MHC-II DCs did not differ (Fig. S7N). Colitis did not drive differences in the abundance of either T-cell subpopulation (Fig. 3N, Fig. SF7O-P). Together, these data demonstrate that systemic inflammation peaked in the colitis model concomitant with peak weight loss and disease severity and may be marked by an elevated presence of myeloid cells, although the inflammation quickly returned to baseline levels after DSS is withdrawn.

### 3.5. Transcriptomic perturbations in the brain coincide with peak disease and exhibit heterogeneity between brain regions

To provide a comprehensive appraisal of the effects of a leaky gut on the brain in the DSS-induced colitis model, we performed high-throughput RNA sequencing of various brain regions, including cortex, hippocampus, ventral midbrain, and striatum from all time points described in Fig. 1A. After discarding genes with low abundance and variability, our dataset covered 21,570 genes. In an initial effort to visualize transcriptomic perturbances during injury and repair in the DSS model, we examined differential expression at each time point relative to the untreated control in the four tissues independently (Fig. 4A). From this analysis, differential expression culminated at the same time as the peaks of disease severity and the systemic immune response. After DSS was withdrawn and mice began to recover, few, if any, genes were differentially expressed in any brain region examined. The exception to this pattern was the striatum; after two weeks of recovery, differential expression increased in the striatum again, with 143 genes significantly regulated compared to 177 genes at 7d of DSS and 149 genes at 2d of recovery.

**Figure 4:**
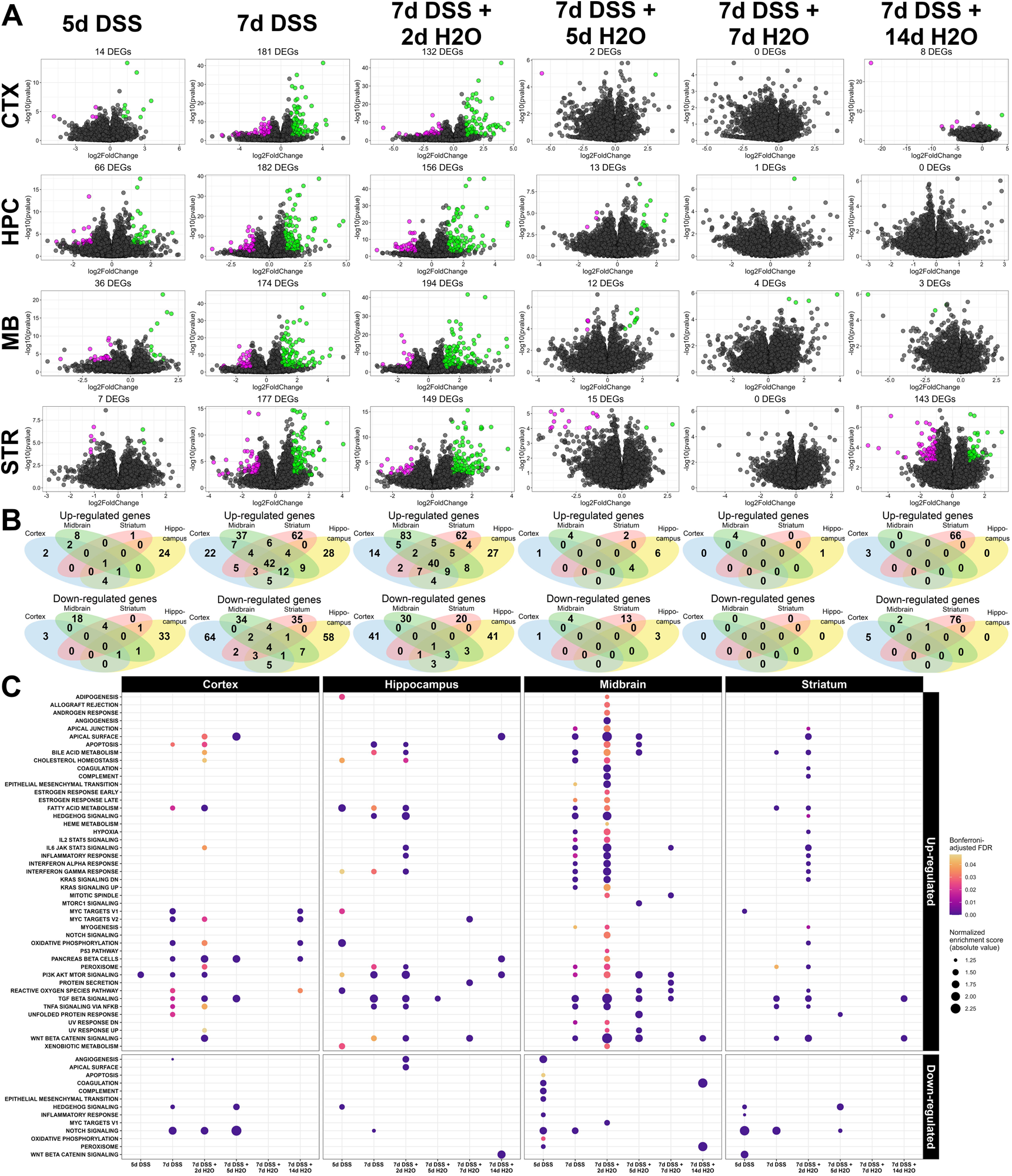
Comparing differential expression in four brain regions during colitis. **A** Volcano plots of all genes in the brain RNA-sequencing datasets, with the log2 transform of the fold change (LFC) from untreated controls on the x-axis and -log10 transform of the p-value on the y-axis. Magenta points are genes with LFC < -1 and Benjamini-Hochberg-adjusted p-value < 0.05, while green points are genes with LFC > 1 and Benjamini-Hochberg-adjusted p-value < 0.05. The total number of differentially expressed genes (adjusted p-value < 0.05) are shown above each panel. **B** Venn diagrams showing how many significantly up-regulated (green genes; top) and significantly down-regulated (magenta genes; bottom) are shared or unique between brain regions (cortex = blue, hippocampus = yellow, midbrain = green, striatum = red) at the time-point described at the top of the column. **C** GSPA output comparing gene lists from each tissue and time point ranked according to LFC after LFC shrinkage. The human MSigDB Hallmark pathways were input as gene sets of interest. Bubbles are colored by a Bonferroni-corrected FDR of enrichment and sized by their normalized enrichment score. GSPA outputs are grouped by tissue (columns) and sign of enrichment score (rows).

To determine whether gene expression profiles differed between brain regions, the lists of up-and down-regulated genes were compared using four-way Venn diagrams at each time point (Fig. 4B). About 39% and 33%, respectively, of up-regulated DEGs at 7d of DSS and 2d of recovery were shared by at least two brain regions, 15-30% of genes were unique to either the midbrain or striatum at both time points and 5-12% of genes were unique to either the cortex or hippocampus. Almost all down-regulated DEGs were unique to their respective brain regions, with at most 14% of genes shared by at least two brain regions. To further compare the expression dynamics in each brain region, we performed a gene set proximity analysis (GSPA) (Cousins et al., 2023) on the set of ranked gene lists from each tissue and time point. This analysis considers the whole interconnectedness of individual genes, employing unsupervised embeddings based on high-confidence protein-protein interactions from the STRING repository (Szklarczyk et al., 2015). As such, this method is more sensitive to implied alterations in a pathway of interest than its precursor gene set enrichment analysis (GSEA). In Fig. 4C, GSPA was applied to all ranked gene lists using the human MSigDB hallmark pathways <u>(Liberzon et al. 2015)</u>. This analysis revealed that the hypoxic response and certain immune signatures, such as interferon signaling, IL-6 signaling, and complement signaling, were preferentially activated in the nigrostriatal system at peak colitis (Fig. 4C). Meanwhile, fatty acid metabolic signatures, TNF signaling via NFkB, TGFß signaling, and Wnt/ß-catenin signaling were activated globally during peak disease, while Notch signaling was suppressed during DSS in all regions to varying degrees (Fig. 4C). Moreover, the striatum did not show evidence for the enrichment of apoptotic signatures, mTOR activation, or cholesterol processing during DSS in contrast to the cortex, hippocampus, and midbrain, while it uniquely re-activated TGFß signaling after two weeks of recovery (Fig. 4C).

In a final effort to compare disparate brain regions in their response to a leaky gut, we built a consensus co-expression network from all four tissues using the WGCNA pipeline (Langfelder and Horvath, 2008). A signed hybrid adjacency network was created using a soft thresholding power of 6. This power was chosen to maximize the scale-free topology model R^2^ in as many tissues as possible (Fig. SF6A-D). Notably, the pattern of connectivity between all regions appeared to differ, with the cortex showing greater connectivity at all soft thresholding powers tested (Fig. SF6A-D). Weighted topological overlap was calculated for each node in each network, and the minimum weight was selected as the consensus overlap. Modules were identified via hierarchical clustering with average linkage, and modules with 75% similarity were merged, leaving 35 distinct consensus co-expression modules (Fig SF6E).

Our consensus network analysis suggested that co-expression patterns were well preserved across the four regions tested here. The mean density of module preservation between pairs of tissues ranged from 0.76 to 0.85, and the patterns of intramodular correlations were similar in all tissues (Fig. SF7), demonstrating overall similar network structures among brain regions. However, many consensus modules were fragmented in set-specific networks (Sig. SF8), suggesting that although the genes in these modules were co-expressed to a degree in every region, the patterns were not identical. The cortex showed overall fewer modules from its set-specific network construction, which agrees with the greater mean connectivity between nodes seen in Fig. SF6A-D, and many consensus modules could be found in one cortex-specific module (Fig. SF8A). In summary, evidence from differential expression analysis, ranked gene list comparisons, and building a consensus co-expression network highlights that patterns of gene adjacency and differential regulation patterns were broadly similar between the cortex, hippocampus, ventral midbrain, and striatum. This similarity motivated us to explore global transcriptomic perturbations in the brain as a result of intestinal inflammation using the consensus co-expression network, which tends to provide more nuanced insight into the biological processes evoked by experimental manipulation.

### 3.6. Transcriptomic perturbations in the brain are associated with colitis severity and systemic inflammation

To begin identifying DSS-associated modules, module eigengenes were computed for all consensus modules and correlated with disease phenotype indexes previously described here (Fig. 1, Fig. 3). Four brain consensus modules showed significant correlations with disease phenotypes in all four regions; the ‘midnightblue’ module was negatively associated with disease severity and circulating markers of inflammation, while the ‘paleturquoise,’ ‘darkgrey,’ and ‘cyan’ modules were positively correlated with these external traits (Fig. 5). To identify brain co-expression modules that may be regulated during different phases of disease, colon consensus modules were categorized into four different types: up-regulated during disease, down-regulated during disease, repair-associated, and persistently regulated (Fig. S11). The ‘midnightblue,’ ‘paleturquoise,’ ‘darkgrey,’ and ‘cyan’ modules showed associations with distal colon modules regulated specifically during DSS administration (Fig. S12). We observed few significant correlations between brain consensus module eigengenes and distal colon eigengenes regulated during the repair phase post-colitis or regulated persistently throughout the dosing scheme (Fig. S12). Additionally, the relationships between consensus module expression and external traits were overall similar between brain regions; we do not observe any brain region-specific associations between co-expression modules and disease phenotypes (Fig. 5, Fig. S12). These analyses provide further support that differential expression in the brain resulting from intestinal permeability was restricted to the height of disease severity. Moreover, we find that the overall gene expression profile was similar between brain regions, as co-expression modules adopt similar profiles during gut leakiness in all four tissues examined here. The four differentially regulated modules highlighted above will be the topic of further dissection.

**Figure 5:**
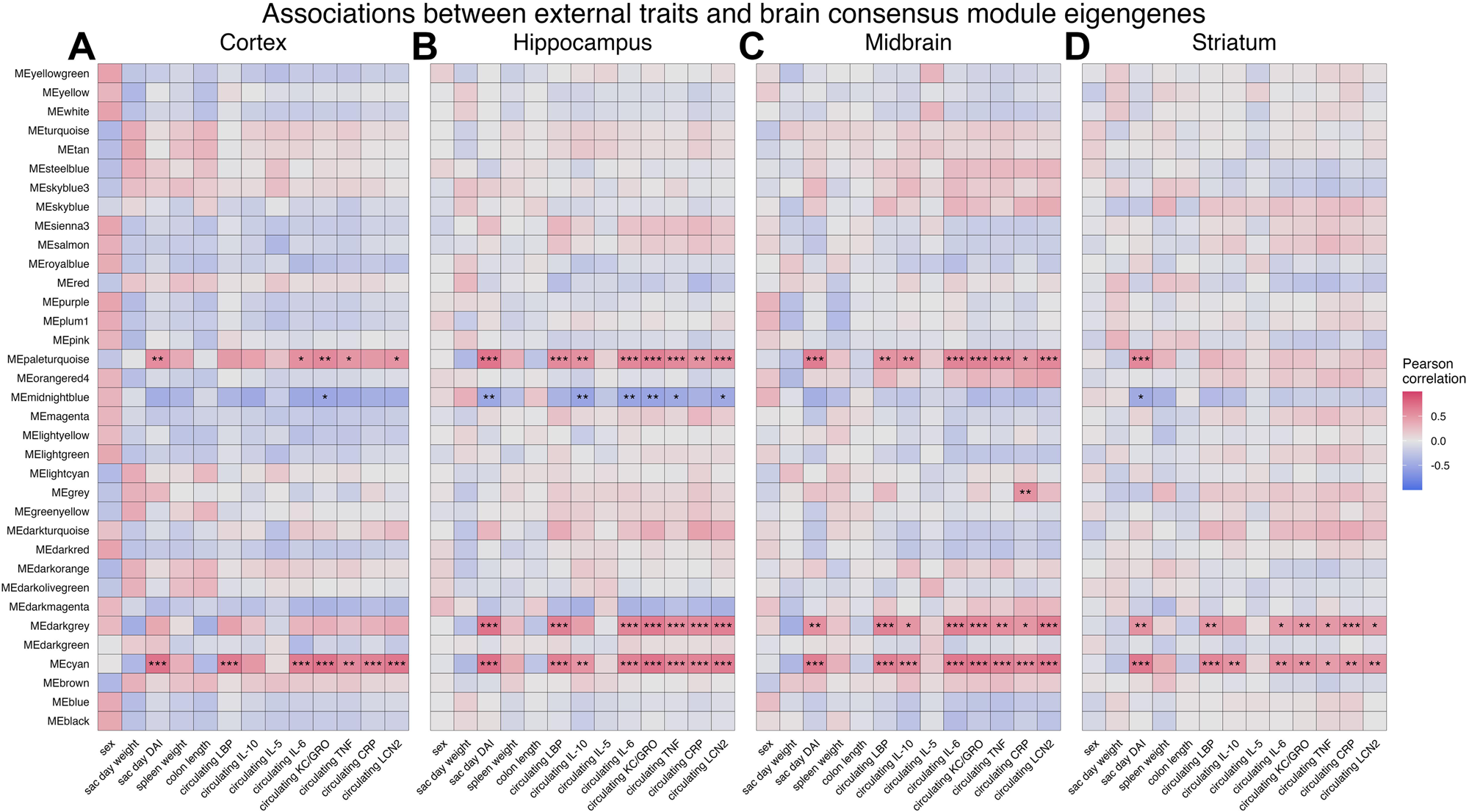
Perturbations in gene expression in the brain are associated with colitis severity and systemic inflammation. **A** Heatmaps of Pearson correlation coefficients between brain consensus module eigengenes and external traits presented in Figure 1 and Figure 3. **B** Heatmap of Pearson correlation coefficients between brain consensus module eigengenes and consensus module eigengenes from the distal colon network, grouped by character and ordered by level of preservation in publicly available datasets. Brain module names are on the y-axis (rows) and text within cells show the p-value of each association. Where all correlations in the four brain regions agreed in sign, the weakest correlation and largest p-value are shown. Where correlations did not agree, cells are gray and blank.

### 3.7. Colitis induces immune activation and oxidative stress response in the brain

We have identified three brain co-expression modules whose expression levels positively relate to disease severity in DSS-induced colitis (Fig. 5). Direct examination of the eigengenes for these modules revealed the expected maximum at 7d of colitis and 2d of recovery when systemic immune markers were high and disease severity was at its worst (Fig. 6A-C). While the overall expression profiles of these modules agreed between regions, differences were notable in the dynamics of individual genes, especially as module membership decreases (Fig. 6D-F). Ranking genes by their meta-module membership identified hub genes for each module, including *Plin4*, *Arrdc2*, *Hif3a*, and *Lcn2* in the ‘cyan’ module (Fig. 6D); *Pnpla2*, *Ly6c1*, *Cdkn1a*, *Ly6a*, and *Fcgr3* in the ‘darkgrey’ module (Fig. 6E); and *Pdk4*, *Arrdc3*, *Tlr7*, and *Dnajc3* in the ‘paleturquoise’ module, which could differ slightly in region-specific module membership (Fig. 6F).

**Figure 6:**
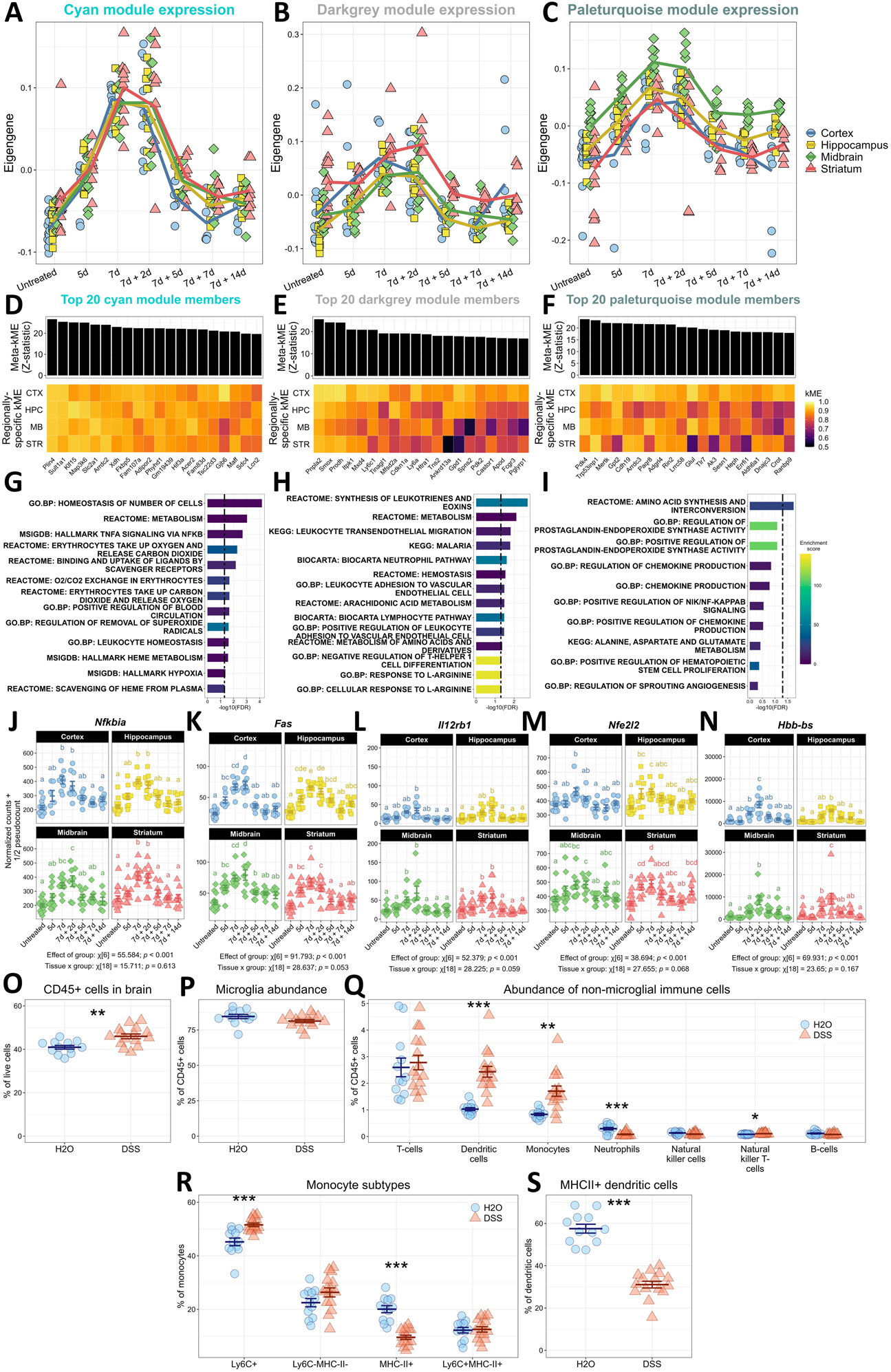
Colitis induces immune activation in the brain. Eigengene plots for the ‘cyan’ module (**A**), ‘darkgrey’ module (**B**), and ‘paleturquoise’ module (**C**), where the line that connects each group represents the mean. Shape/color distinguish eigengenes from different regions according to the legend on the right. Bar charts showing the meta-module membership and corresponding regionally-specific module membership (i.e., correlation with the module eigengene, kME) in a heatmap below for the ‘cyan’ (**D**), ‘darkgrey’ (**E**), and ‘paleturquoise’ (**F**) modules. In these three figures, genes were sorted according to their meta-module membership and the top 20 members are shown for each. Gene set enrichment results for the ‘cyan’ (**G**), ‘darkgrey’ (**H**), and ‘paleturquoise’ (**I**) module, ranked according to their unadjusted *p*-value. Bars represent the -log10 transform of the FDR, and the dashed line shows where FDR < 0.05. Bars are colored according to their enrichment score, which is a measure of how much of the gene set is found in the module versus how much is found in the background. DESeq2-normalized counts for *Nfkbia* (**J**), *Fas* (**K**), *Il12rb1* (**L**), *Nfe2l2* (**M**), and *Hbb-bs* (**N**) are shown. Statistical results are shown with the plots, gathered from the multiple mixed-effects modeling procedure described in the Methods section. *Post hoc* comparisons were made within each region, where groups that share a letter are not statistically significantly different (*p* > 0.05). **O** Percentage of live cells in the brain that are CD45+ are DSS, according to the experimental details in Fig. 3 (Student’s *t*[25] = 3.475, *p* = 0.002). **P** Percentage of CD45+ cells that are microglia. **Q** Abundances of non-microglial immune cells in the brain shown as a percentage of CD45+ cells (two-way ANOVA: main effect of DSS – *F*[1, 25] = 0.468, *p* = 0.500; main effect of population – *F*[1.104, 27.604] = 6858.273, *p* < 0.001; interaction – *F*[1.104, 27.604] = 3.844, *p* = 0.056). *Post hoc* testing was conducted within each cell type and *p*-values were adjusted with Bonferroni’s correction. The two-way ANOVA was conducted using data from **P** and **Q** as the relative abundance of each population is not independent from each other. **R** Relative abundances of monocyte subtypes in the brain (two-way ANOVA: main effect of DSS – *F*[1, 25] = 0, *p* = 1; main effect of population – *F*[2.091, 52.267] = 274.041, *p* < 0.001; interaction – *F*[2.091, 52.267] = 14.024, *p* < 0.001). *Post hoc* testing was conducted within each cell type and *p*-values were adjusted with Bonferroni’s correction. **S** Percentage of dendritic cells in the brain that are MHC-II+ (Student’s *t*[25] = -10.527, *p* < 0.001). All data are shown as mean ± SEM. \*\**p* < 0.05, \*\**p* < 0.01, \*\*\**p* < 0.001.

A gene set enrichment analysis revealed enrichment of pathways associated with immunological processes in all three up-regulated modules (Fig. 6G-I), which is consistent with many prior studies (Craig et al., 2022). These pathways include leukocyte activation and TNF signaling via the NF-κB inflammasome in the ‘cyan’ and ‘darkgrey’ modules. Chemokine production pathways were found to be enriched in the ‘paleturquoise’ module, but its enrichment did not survive FDR correction, likely due to the small size of the module in general. Examination of individual genes revealed a complex immune activation in the brain due to colitis. Some genes were regulated globally during colitis, including *Nfkbia*, *Fas*, and *Il12rb1*, showing no significant interaction between colitis and region (Fig. 6J-L). Other genes were regulated in a tissue-specific manner (i.e., showed a significant interaction between region and DSS), including *Lyve1*, *C4a*, and *Cd14* (Fig. S13). Our co-expression analysis also revealed hitherto unrealized pathways induced in the brain by a leaky gut. Oxidative stress response was enriched in the ‘cyan’ module, marked by the global regulation of *Nfe2l2* during colitis independent of brain region (Fig. 6M). The ‘cyan’ module showed the enrichment of pathways related to oxygen levels and heme metabolism, stemming from the global up-regulation of several hemoglobin genes, including *Hbb-bs* (Fig. 6N), *Hba-a1*, and *Hbb-bt* (Fig. S13). Several other genes implicated in these pathways are shown in Fig. S13 and summarized in Supplemental File 4. Gene-level information for every gene included in our dataset, including module membership statistics and gene-trait relationships, can be found in Supplemental File 2. Unlike several other published reports (Batra et al., 2021; He et al., 2021; Sroor et al., 2019; Talley et al., 2021; Vitali et al., 2022), we did not observe a significant up-regulation of genes encoding canonical pro-inflammatory cytokines *Tnf* (Fig. S13JJ) or *Il1b* (Fig. S13KK) or a key chemoattractant *Ccl2* (Fig. S13LL) in the brain during our DSS dosing schedule.

To validate the enrichment of leukocyte migration and adhesion systems, we performed flow cytometry to characterize immune cell populations in the brain during peak colitis, as shown in Fig. 3H. In accordance with the data above, a greater proportion of the cells in the brain during colitis were CD45+ (Fig. 6O, Fig. S14B). There were no differences in the relative abundance (Fig. 6P) or count of microglia (Fig. S14C) after DSS, consistent with a previous study (Sroor et al., 2019), nor were there differing amounts of MHC-II+ microglia (Fig. S14D-E). However, there were greater abundances of dendritic cells (Fig. 6Q, Fig. S14F), monocytes (Fig. 6Q, Fig. S14G) and natural killer T-cells (Fig. 6Q, Fig. S14H) in the brain with a leaky gut. More monocytes were Ly6C+ (Fig. 6R, Fig. S14I) while fewer of the monocytes were MHC-II+ (Fig. 6R, Fig. S14J) in the brain with DSS, and we observed no differences evoked by DSS in the relative abundance of Ly6C-MHC-II-monocytes (Fig. 6R) or Ly6C+MHC-II+ monocytes (Fig. 6R) although these two subtypes displayed greater raw counts with colitis (Fig. S14K-L). MHC-II+ dendritic cells displayed higher raw counts in the DSS brain (Fig. S14M) although a smaller proportion of total DCs were MHC-II+ (Fig. 6S). Meanwhile, there was a lesser abundance of neutrophils in the brain according to their overall counts (Fig. S14N) and relative frequency (Fig. 6Q). There were no changes in the presence of T-cells (Fig. 6Q, Fig. S14O), natural killer cells (Fig. 6Q; Fig. S14P), or B-cells (Fig. 6Q; Fig. S14Q) in the brain. Additionally, there appeared to be no changes in the presence of T-cell subtypes in the brain with colitis, reflected by no differences in the percentages or counts of T-cells that are CD4+ (Fig. S14R-S) with a poor detection of CD8+ T-cells (Fig. S14T-U). These data demonstrate that peak colitis drove global although transient immune activation in the brain, marked by transcriptomic signatures associated with leukocyte recruitment and NF-κB inflammasome signaling. Notably, we observed the recruitment of myeloid cells, namely dendritic cells and monocytes, to the brain, implicating a key role for the innate immune system in leaky gut-induced brain injury or dysfunction.

### 3.8. A leaky gut induces transient loss of myelin support in the brain

Consensus co-expression network analysis highlights the ‘midnightblue’ module as the only co-expression module that was consistently negatively associated with disease markers and circulating inflammatory molecules in most brain regions (Fig. 5). This module showed a decrease in the eigengene during the DSS dosing paradigm with a return to baseline levels thereafter (Fig. 7A). Notably, this decrease in expression was similar across regions despite the baseline differences between them. Several of this module’s strongest gene members are notable oligodendrocyte genes, including *Gjb1*, *Mag*, *Sox10*, *Gjc2*, *Opalin*, and *Cnp* (Fig. 7B). Accordingly, gene set enrichment analysis revealed an enrichment of pathways associated with myelination and oligodendrocyte proliferation/development (Fig. 7C, Supplemental File 4). Normalized count data are shown for many of these genes in Fig. 7D-M. Gene-level statistics including gene-trait relationships and module membership data can be found in Supplemental File 2. In many cases, the effect of DSS was not obvious in separate tissues, although mixed-effects modeling suggests that DSS significantly affects the expression of these genes. Our unbiased network approach demonstrates that the dysregulation of oligodendrocyte function or myelin maintenance was a global consequence of a leaky gut.

**Figure 7:**
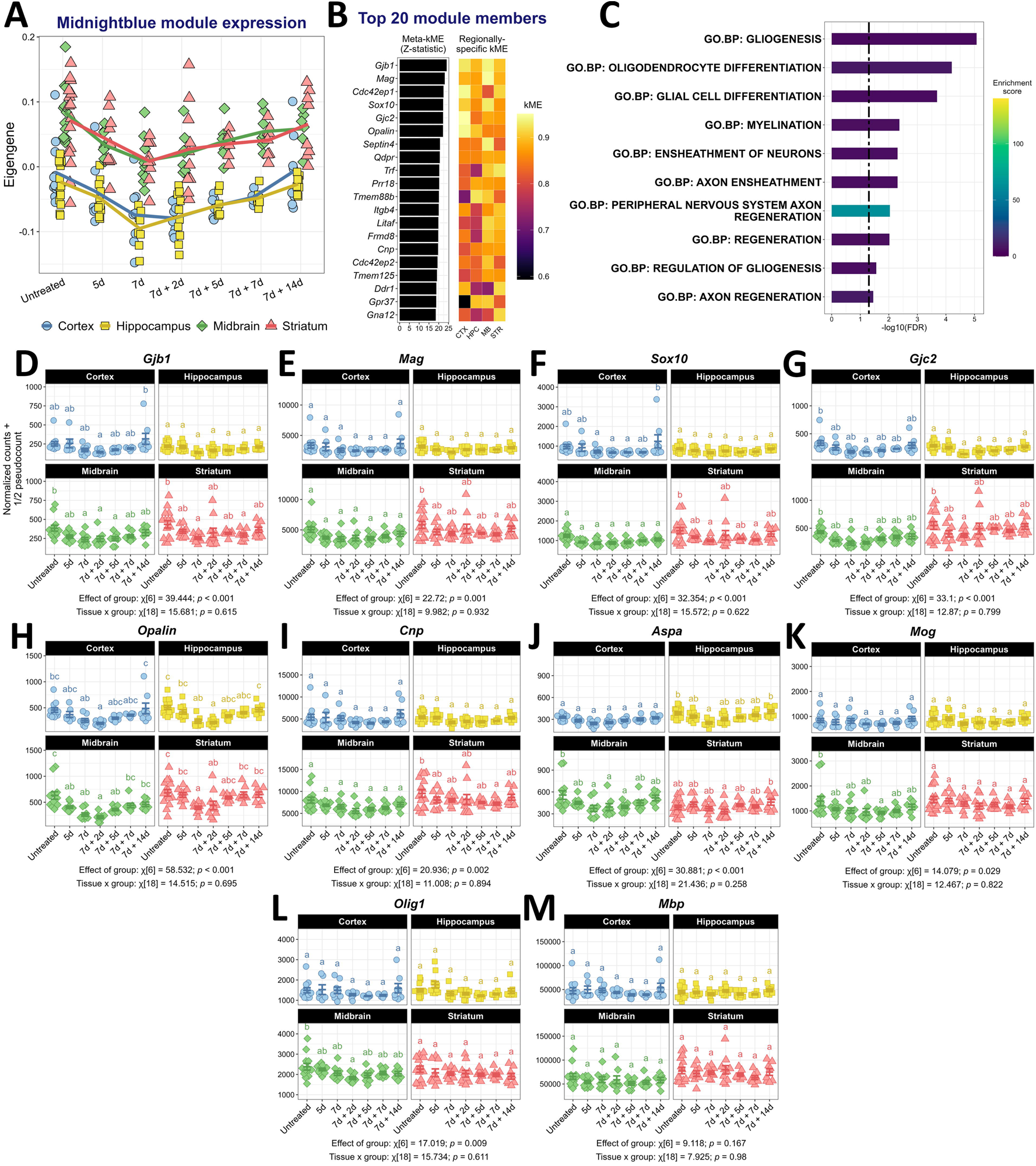
A leaky gut induces transient loss of myelin support in the brain. **A** Eigengene of the ‘midnight’ blue module shown over time in each brain region, distinguished by shape and color. The line connecting each group represents the mean. **B** Gene meta-module membership statistics with the corresponding regionally-specific module membership for the top 20 genes in the ‘midnightblue’ module, ranked by their meta-module membership. **C** Gene set enrichment results, with gene sets ranked by their unadjusted *p*-value. Bars represent the -log10 transform of the FDR and are colored by their gene set’s enrichment score. Bars reaching to the right of dashed line have an FDR < 0.05. DESeq2-normalized counts for myelin genes assigned to the ‘midnightblue’ module, including *Gjb1* (**D**), *Mag* (**E**), *Sox10* (**F**), *Gjc2* (**G**), *Opalin* (**H**), *Cnp* (**I**), *Aspa* (**J**), *Mog* (**K**), *Olig1* (**L**), and *Mbp* (**M**). Statistical results are shown with the plots, gathered from the multiple mixed-effects modeling procedure described in the Methods section. *Post hoc* comparisons were made within each region, where groups that share a letter are not statistically significantly different (*p* > 0.05). Data are shown as mean ± SEM.

## 4. Discussion

The present study offers an unbiased appraisal of disturbances along the gut-brain axis in a ubiquitous model of experimental colitis. Although we expected that a comparison of outcomes between our study and previously published ones may reflect differences between experimental sites, including pathogen load in animal facilities and DSS reagent vendors and lots, we find that repair mechanisms post-injury in the DSS model are highly conserved between studies. Also, our results suggest that gut immune activation, metabolic perturbation, and stress responses are broadly similar between studies. Our findings implicate predominately innate immune involvement in the intestine, a well-established feature of the model (Kiesler et al., 2015). Our findings also support the generalizability of the DSS model, bolstering its usefulness in modeling human disease. A recent study found success in stratifying UC patients based on DSS data to identify non-responders to medication (Czarnewski et al., 2019). However, we report study-specific responses, including a uniquely persistent and interferon-lacking immune response in our hands, a unique activation rather than suppression of metabolic gene programs in GSE131032, and varied degrees of network-level similarity between studies. Future efforts will attempt to identify the factors that drive these discrepancies.

Here, we focused on the circulating immune system as a conduit between the gut and the brain (Houser and Tansey, 2017). We found peak colitis-associated increases in circulating LBP, CRP, TNF, IL-10, and KC/GRO, likely from the mobilization of myeloid cells. This humoral profile overlaps with that of human IBD, which shows similar levels of circulating cytokines, endotoxemia markers, and innate immune cells (Korolkova et al., 2015; Kwon et al., 2022; Okba et al., 2019), further upholding the translational value of the DSS model. Importantly, the circulating profile may help explain neuropsychiatric comorbidities associated with IBD; circulating cytokines, including those measured here, are also raised in depressive disorders (Berk et al., 2013; Gałecki et al., 2012; Pasco et al., 2010), anxiety disorders (Costello et al., 2019; Vogelzangs et al., 2013), ASD (Enstrom et al., 2009; Onore et al., 2012; Yin et al., 2020), MS (Bai et al., 2019), and PD (Tansey et al., 2022). Circulating immune markers also correlate with emotional and cognitive symptoms in IBD (Abautret-Daly et al., 2017; Iordache et al., 2022; Nemirovsky et al., 2022), leading the gut-brain propagation of immune signals via the circulation to be a prime suspect in the induction of neuropsychiatric disorder with intestinal disease (Craig et al., 2022). Indeed, several studies employing the DSS model suggest that brain effects depend on a systemic response (Jackson et al., 2022; Maltz et al., 2022), and the relationship between systemic inflammation and brain transcriptomic changes in our study corroborates this.

Our transcriptomic approach supports the field’s current emphasis on inflammation as the principal consequence of a leaky gut on the brain. Our data support the role of the S100 alarmin S100A9 (Talley et al., 2021) and implicate innate immune cells rather than microglia (Batra et al., 2021; Gampierakis et al., 2021; Sroor et al., 2019). Our data also provide evidence for TNF signaling as a key system in gut-induced neuroinflammation due to the up-regulation of several downstream signaling molecules, including *Nfkbia*, *Icosl*, *Bhlhe40*, and *Fosb* (Gampierakis et al., 2021; Han et al., 2018; Lin et al., 2021; Talley et al., 2021) even though we do not find *Tnf* itself to be up-regulated with colitis. We observed a transient neuroimmune activation, which contrasts our hypothesis that DSS would trigger persistent neuroinflammation. A previous study from our laboratory achieved lasting neuroinflammation with DSS after a two-hit model of PD, marked by T-cell infiltration and interferon signaling after a single bout of colitis, neither of which we can corroborate here, emphasizing a need for multiple genetic and environmental sources of vulnerability to elicit such a robust response (Houser et al., 2021; Kline et al., 2021). In general, this hypothesis is supported by several studies that demonstrate lasting neuroinflammation or overt brain injury with DSS only when paired with other insults, such as aging, genetic mutations affecting the immune system, or a secondary disease model, and may require several dosing rounds to achieve chronicity (Cabezudo et al., 2023; Feng et al., 2019; Gampierakis et al., 2021; He et al., 2021; Kishimoto et al., 2019; Lin et al., 2021; Zonis et al., 2015).

Our approach also reveals a potential effect on brain myelination during colitis. A recent report demonstrates a similar effect of DSS using immunoblotting (Takahashi et al., 2023), lending to the robustness of this effect. This brain effect was accompanied by depressive behaviors and a loss in serotonergic tone, suggesting that demyelination may be a physiological basis for psychiatric comorbidities in IBD. Although, it is presently unknown whether IBD patients exhibit either brain demyelination or serotonin deficits.

Our approach also implicates novel molecular processes are disrupted in the brain in response to increased intestinal permeability. We report a global overexpression of hemoglobin genes, the hypoxia-induced transcription factor *Hif3a*, and *Nfe2l2*, the gene encoding for the Nrf2. Together, this collection of genes may reflect oxidative stress in neurons, potentially in response to altered oxygen levels in the brain (Heidbreder et al., 2003; He et al., 2020, 2010; Sarlus et al., 2013; Schelshorn et al., 2009). It would be valuable to validate and extend these findings, including the measurement and localization of reactive oxygen species and the direct examination of hypoxia in the brain during colitis.

Discussion around the gut-brain axis would be incomplete without mention of the intestinal microbiome, a diverse flora with many recorded interactions with the immune and central nervous systems. It is well documented that DSS-induced colitis results in dysbiosis, with a loss in anti-inflammatory microbes and short-chain fatty acid producers (Lee et al., 2023; Munyaka et al., 2016), which may mimic the dysbiosis in IBD (Caruso et al., 2020). Dysbiosis has also been observed in ASD (Fowlie et al., 2018), PD (Scheperjans et al., 2015; Wallen et al., 2022; Xie et al., 2022), MS (Chen et al., 2016; Jangi et al., 2016) and depressive and anxiety disorders (Dinan and Cryan, 2019; Radjabzadeh et al., 2022; Wan et al., 2022), demonstrating another potential link between gut and brain. Accordingly, some rodent studies using DSS implicate the microbiome as the cause of behavioral symptoms rather than systemic immunity (Chen et al., 2023; Salvo et al., 2020; Vicentini et al., 2022; H. Zhao et al., 2023; L.-P. Zhao et al., 2023), although the nature of this microbiome-brain interaction is mysterious. The translocation of bacteria or metabolites from gut to circulation is documented in DSS (Kwon et al., 2021), IBD (Caradonna et al., 2000), and neuropsychiatric diseases (Emanuele et al., 2010; Forsyth et al., 2011), but a gut-to-brain translocation of bacteria has not been reported and may be a fruitful line of investigation.

We suspect the immune system and microbiome are more likely to interact in this model and human disease, so consideration for both must be given. Altering the microbiome in the DSS model affects colitis itself, which likely affects the leakage of microbial products and inflammatory mediators from the gut to circulation in turn (Lee et al., 2022; Lu et al., 2022; Peng et al., 2020; Qu et al., 2021). Additionally, microbiota-T-cell interactions have been reported that induce demyelination (Lynch et al., 2021; Merchak et al., 2023), and gut-educated T- and B-cells have been found in the meninges after colitis (Feng et al., 2019; Fitzpatrick et al., 2020) which may in turn affect the brain’s innate immunity (Hobson et al., 2023). Our data suggest a predominately innate immune response in naïve, wild-type, young mice with colitis, while other studies emphasize the mobilization of these adaptive immune cells amidst other experimental insults. As such, we find it likely that the development of neuropsychiatric comorbidities in IBD and the involvement of sub-clinical gastrointestinal distress in neuropsychiatric disorders may depend on other intervening variables, including the state of the gut flora. Certainly, consideration of intestinal health and flora has helped stratify patients with PD (Wijeyekoon et al., 2020) and ASD (Chen et al., 2022; Rose et al., 2018) into clinically distinct subtypes, which may inform future therapeutic avenues in a more personalized manner.

Intestinal permeability and inflammation have become increasingly important features of disorders long thought to be limited to the brain (Berk et al., 2013; Houser and Tansey, 2017; Morton et al., 2023), and the brain has become increasingly implicated in IBDs (Bernstein et al., 2019; Günther et al., 2021). Our study is the first to apply an unsupervised analysis to the gut-brain axis of a prevalent colitis model, which supports innate immune activation and demyelination as principal components of the brain’s response to a leaky gut. We suggest that these effects are associated with disease severity and systemic inflammation. Moreover, we examine the reproducibility of many central disease processes in the gut of the DSS-induced colitis model, which may serve as a foundation for further examination of the gut-to-brain propagation of injury in this model. Future endeavors will examine microbiome-immune interactions present in the DSS model as well as human diseases marked by gut leakiness, focusing on genetic and environmental factors that sway the microbiome and/or the immune system. Determining the cause of brain inflammation and injury during intestinal permeability will yield therapeutic insight into multi-system mechanisms of various human neurological diseases.

**Supp. Table 1: Disease activity index (DAI) scoring rubric.**

**Supp. Table 2: Product and use information for all antibodies used in the present study.**

**Supp. Table 3: Primer sequences used for qPCR.**

**Supp. Figure 1: Consensus co-expression network construction in two colon segments. A** Scale-free topology model R^2^ in distal (green) and proximal (pink) colon segment RNA sequencing datasets as a function of soft thresholding/beta power. The median (**B**), average (**C**), and maximum (**D**) connectivity of both segments are shown as functions of soft thresholding power. From these data, a soft power of 7 was selected to reach a model fit above the recommended threshold of 0.8, which is also the point at which connectivity does not decrease substantially as soft threshold power increases. **E** Gene dendrogram with original (unmerged) and merged module labels. Modules were merged based on average linkage distance shown in the dendrogram in **F**.

**Supp. Figure 2: Assessment of gene network preservation between distal and proximal colon segments. A** Intermodular correlation heatmap in distal colon. Each row and column correspond to the eigengene of the module. Red represents a positive correlation between modules, while blue represents a negative correlation. **B** Bar chart showing the degree of preservation of each consensus module in both colon networks, where taller bars indicate a greater degree of preservation. The overall high D-value shown at the top indicates a strong preservation of modules between both networks. **C** Heatmap displaying the difference between networks in eigengene adjacencies. Preservation is calculated as one minus the absolute value of the difference of the eigengene correlations in the two colon networks. Thus, a brighter red indicates a smaller difference between networks. **D** Intermodular correlation heatmap in proximal colon, coded the same as **A**. **E** Multiple Fisher’s exact test comparing the independence of gene assignment in distal colon-specific modules and gene assignment to consensus modules. Rows are consensus modules and columns are distal colon-specific modules. Cells shaded red indicate a significant Fisher’s exact test (*p* < 0.05) after Bonferroni’s correction, meaning there is significant overlap between the distal colon-specific module and the consensus module. The number in each cell reflects the number of genes overlapping the two modules shown at that intersection. **F** Multiple Fisher’s exact test comparing the independence of gene assignment in proximal colon-specific modules and gene assignment to consensus modules, coded the same as in **E**.

**Supp. Figure 3: Heatmap showing the degree of preservation of colon consensus modules in three publicly available datasets.** The x-axis describes the reference-test pair of datasets used for that respective column. The reference dataset is one of our two colon segments, where the connectivity profile from that dataset was evaluated for preservation in the test dataset. The y-axis names modules, which are grouped according to their inferred function. In the test dataset, gene module assignment was randomly permuted 200 times, and the mean and standard deviation from these permutations was used to convert connectivity statistics from the actual module assignment to a Z-score. A Z-score between 2 and 10 indicates “weak-to-moderate” support for the preservation of that module in the test network based on the profile of the reference network, while a Z-score above 10 indicates “strong” support.

**Supp. Figure 4: Reproducible and experiment-specific stress responses in the intestine during DSS. A-D** Gene set enrichment results from the ‘coral3,’ ‘coral2,’ ‘white,’ and ‘thistle2’ colon consensus modules. The top 10 pathways are shown, ranked on their unadjusted *p-*values. Bars show the -log10 transform of the FDR for that gene set. Bars reaching to the right of the dashed line reflect an FDR < 0.05. **E-H** Module eigengenes for each of four datasets: top-left shows our two datasets colored by colon segment; top-right shows GSE131032; bottom-left are the eigengenes from GSE168053 colored by colon segment; bottom-right shows GSE210405. The x-axis indicates day of the study, and the red shading indicates when DSS was administered. **I-L** Meta-module membership shown as a Z-statistic based on our two colon datasets with the corresponding dataset-specific module membership (correlation with the module eigengene, kME) for each gene displayed in a heatmap. The top 30 module members, defined by their high meta-kME, are shown here for each module.

**Supp. Figure 5: Reproducible and study-specific metabolic disturbances in the colon during colitis. A-G** Gene set enrichment results from the ‘brown,’ ‘coral,’ ‘darkolvegreen4,’ ‘lightpink3,’ ‘coral1,’ ‘indianred4,’ and ‘darkslateblue’ colon consensus modules. Gene sets are ranked by ascending *p-*value, and the top 10 pathways are displayed. Bars show the -log10 transform of the FDR for that gene set, and bars rising to the right of the dashed line reflect an FDR < 0.05. **H-N** Module eigengenes for each of four datasets: top-left shows our two datasets colored by colon segment; top-right shows GSE131032; bottom-left are the eigengenes from GSE168053 colored by colon segment; bottom-right shows GSE210405. The x-axis indicates day of the study, and the red shading indicates when DSS was administered. **O-U** Meta-module membership shown as a Z-statistic based on our two colon datasets with the corresponding dataset-specific module membership (correlation with the module eigengene, kME) for each gene displayed in a heatmap. The top 30 module members, defined by their high meta-kME, are shown here for each module.

**Supp. Figure 6: Reproducible and study-specific intestinal repair processes during and after DSS-induced colitis. A-D** Gene set enrichment results from the ‘floralwhite,’ ‘lightcoral,’ ‘salmon,’ and ‘tan’ colon consensus modules. The top 10 pathways are shown, ranked on their *p-*values. Bars show the -log10 transform of the FDR for that gene set, and bars rising to the right of the dashed line have an FDR < 0.05. **E-H** Module eigengenes for each of four datasets: top-left shows our two datasets colored by colon segment; top-right shows GSE131032; bottom-left are the eigengenes from GSE168053 colored by colon segment; bottom-right shows GSE210405. The x-axis indicates day of the study, and the red shading indicates when DSS was administered. **I-L** Meta-module membership shown as a Z-statistic based on our two colon datasets with the corresponding dataset-specific module membership (correlation with the module eigengene, kME) for each gene displayed in a heatmap. The top 30 module members, defined by their high meta-kME, are shown here for each module.

**Supp. Figure 7: Identification of peripheral immune cells with multi-color flow cytometry. A** Gating strategy to identify circulating immune cells with flow cytometry. **B** Raw counts of CD45+ cells isolated from blood (Student’s *t*[25] = -0.530, *p* = 0.601). **C** Raw counts of circulating B-cells (Student’s *t*[25] = -4.000, *p* < 0.001). **D** Raw counts of circulating monocytes (Welch’s *t*[14.605] = 3.871, *p* = 0.002). **E** Raw counts of circulating dendritic cells (Welch’s *t*[16.692] = 4.671, *p* < 0.001). **F** Raw counts of neutrophils in circulation (Welch’s *t*[17.461] = 0.976, *p* = 0.342). **G** Counts of total T-cells in circulation (Student’s t[25] = -0.955, *p* = 0.351). **H** Counts of natural killer cells in circulation (Mann-Whitney U = 78, *p* = 0.575). **I** Counts of circulating natural killer T-cells (Mann-Whitney *U* = 39, *p* = 0.014). **J** Counts of Ly6C-high monocytes (Welch’s *t*[14.666] = 3.405, *p* = 0.004). **K** Counts of Ly6C-low monocytes (Welch’s *t*[14.496] = 3.714, *p* = 0.002). **L** Counts of MHC-II+ monocytes (Welch’s t[16.333] = 1.595, *p* = 0.130). **M** Counts of Ly6C+MHC-II+ monocytes (Welch’s *t*[16.604] = 1.742, *p* = 0.100). **N** Counts of MHC-II+ dendritic cells (Welch’s *t*[14.929] = 1.905, *p* = 0.076). **O** CD4+ T-cell counts (Student’s *t*[25] = -0.649, *p* = 0.522). **P** CD8+ T-cell counts (Student’s t[25] = -1.085, *p* = 0.288). Data in **B-P** are presented as mean ± SEM. \**p* < 0.05, \*\**p* < 0.01, \*\*\**p* < 0.001.

**Supp. Figure 8: Consensus co-expression network construction in four brain regions. A** Scale-free topology model R^2^ in cortex (red), midbrain (blue), hippocampus (green), and striatum (yellow) RNA sequencing datasets as a function of soft thresholding/beta power. The median (**B**), average (**C**), and maximum (**D**) connectivity of both segments are shown as functions of soft thresholding power. From these data, a soft power of 6 was selected to maximize the model fit, which was used as the primary criterion of soft power selection due to the discrepancies in connectivity profiles between regions. For example, notice that the cortex dataset displays much higher connectivity than the other three regions at all soft powers examined. **E** Gene dendrogram with original (unmerged) and merged module labels. Modules were merged based on average linkage distance shown in the dendrogram in **F**.

**Supp. Figure 9: Assessment of the preservation of eigengene adjacency between cortex, midbrain, hippocampus, and striatum. A** Intermodular correlation heatmap in cortex samples. Each row and column correspond to the eigengene of a module. Red represents a positive correlation, while blue represents a negative correlation. Bar chart showing the aggregate preservation of individual eigengenes between cortex and midbrain samples (**B**), between cortex and hippocampus (**C**), and between cortex and striatum (**D**). **E** Heatmap displaying the pairwise preservation of modules, defined as one minus the absolute value of the difference in the eigengene adjacencies, in cortex and midbrain samples. **F** Intermodular correlation heatmap in midbrain samples, coded as in **A**. Bar chart showing the aggregate preservation of individual eigengenes between midbrain and hippocampus (**G**) and between midbrain and striatum (**H**) samples. Heatmap displaying the pairwise preservation of modules between hippocampus and cortex (**I**) and between hippocampus and midbrain (**J**), coded as in **E**. **K** Intermodular correlation heatmap in hippocampus samples, coded as in **A**. **L** Bar chart showing the aggregate preservation of individual eigengenes in hippocampus and striatum samples. Heatmap showing the pairwise preservation of modules between cortex and striatum (**M**), between midbrain and striatum (**N**), and between hippocampus and striatum (**O**), coded as in **E**. **P** Intermodular correlation heatmap in striatum samples, coded as in **A**. In short, the diagonal of this matrix shows intermodular correlations in region-specific datasets. Red indicates a positive correlation between eigengenes while blue indicates negative. The upper triangle contains bar charts of eigengene preservation between pairs of datasets and the lower triangle shows heatmaps of the pairwise preservation networks, where “preservation” is calculated as one minus the absolute value of the difference of the eigengene correlations between the two networks. A preservation heatmap or bar chart in row *i* and column *j* show pairwise preservation statistics from sets *i* and *j*, which are named above the eigengene adjacency heatmaps on the diagonal.

**Supp. Figure 10: Assessment of the preservation of gene module assignment in cortex, hippocampus, midbrain, and striatum.** Multiple Fisher’s exact test examining the independence of gene assignment to consensus modules and cortex-specific modules (**A**), hippocampus-specific modules (**B**), midbrain-specific modules (**C**), and striatum-specific modules (**D**). Columns are consensus modules and rows are region-specific modules. Cells shaded red indicate a significant Fisher’s exact test (*p* < 0.05) after Bonferroni’s correction, meaning there is significant overlap between the region-specific module and the consensus module. The number in each cell reflects the number of genes overlapping the two modules shown at that intersection.

**Supp. Figure 11: Classification of colon consensus modules based on eigengenes.** Eigengene plots for every colon consensus module identified by our analysis. The color name of the module is shown in the black title box, and the number of genes assigned to that module is shown in parentheses next to it. Distal colon sample eigengenes are shown by the green circles, and proximal colon eigengenes are shown by the pink squares. Module categorization is shown by the colored boxes around the title box: orange corresponds to modules up-regulated during disease, magenta corresponds to modules down-regulated during disease, green corresponds to modules regulated during the repair phase, and cyan corresponds to modules that appear to be persistently regulated throughout the dosing scheme.

**Supp. Figure 12: Transcriptomic disturbances in brain during colitis are associated with disease-associated gene programs in the intestine.** Heatmaps showing the Pearson correlation coefficient between colon consensus module eigengenes in distal colon (rows) and brain consensus module eigengenes in cortex (**A**), hippocampus (**B**), midbrain (**C**), and striatum (**D**). Red indicates a positive correlation while blue indicates a negative correlation. Numbers in each cell reflect the *p*-value of the correlation after Bonferroni’s correction, rounded to two digits. Distal colon module eigengenes are separated according to their expression profiles outlined in Fig. SF2 and Fig. 2. Note the number of significant correlations between the brain ‘paleturquoise’, ‘midnightblue’, ‘cyan’, and ‘darkgrey’ modules with distal colon modules that are differentially regulated preferentially during DSS administration compared to the paucity of significant correlations between those four modules with distal colon modules that are differentially regulated during the repair phase or consistently throughout the paradigm.

**Supp. Figure 13: DSS-associated differential expression of immune system genes in the brain.** DESeq2-normalized counts for genes related to immune system processes are shown in **A-GG**. Separate facets were created for each brain region, where blue circles correspond to cortex samples, yellow squares correspond to hippocampus samples, green diamonds correspond to midbrain samples, and red triangles correspond to striatum samples. Statistical results are shown with the plots, gathered from the multiple mixed-effects modeling procedure described in the Methods section. Pairwise comparisons were made within each region, and the compact letter display (CLD) is shown. Groups that share a letter within each facet are not statistically significantly different (*p* > 0.05 after Tukey’s correction) from each other. The expression of *Tnf* (one-way ANOVA: F[5, 45] = 2.290, *p* = 0.062) (**HH**), *Il1b* (one-way ANOVA: F[5, 48] = 1.137, *p* = 0.354) (**II**), and *Ccl2* (one-way ANOVA: Welch’s F[5, 19.652] = 1.294, *p* = 0.306) (**JJ**) was measured with qPCR in midbrain samples from all groups excluding the 7d DSS + 14d H2O group. All data in this figure are shown as mean ± SEM.

**Supp. Figure 14: Identification of microglia and other immune cell populations in brain by multi-color flow cytometry. A** The gating strategy to identify microglia and other immune cell populations in the brain with flow cytometry. **B** Raw counts of CD45+ cells in the brain (Student’s *t*[25] = 3.522, *p* = 0.002). **C** Raw counts of microglial cells in the brain (Student’s *t*[25] = 1.683, *p* = 0.105). **D** MHC-II+ microglia abundance expressed as a percentage of total microglia (Student’s *t*[25] = -0.840, *p* = 0.409). **E** Raw counts of MHC-II+ microglia (Student’s *t*[25] = 0.102, *p* = 0.919). **F** Counts of dendritic cells in brain (Welch’s t[15.317] = 7.341, *p* < 0.001). **G** Total monocyte counts in brain (Welch’s *t*[15.050] = 4.761, *p* < 0.001). **H** Natural killer T-cell counts in brain (Student’s *t*[25] = 4.117, *p* < 0.001). **I** Ly6C+ monocyte subtype counts (Welch’s *t*[15.172], *p* < 0.001). **J** MHC-II+ monocyte subset counts (Welch’s *t*[23.256] = 0.370, *p* = 0.715). **K** Counts of Ly6C-MHC-II-monocytes in brain (Welch’s *t*[16.007] = 4.361, *p* < 0.001). **L** Counts of Ly6C+MHC-II+ double-positive monocytes in brain (Welch’s *t*[15.802] = 3.981, *p* = 0.001). **M** Counts of MHC-II+ dendritic cells (). **N** Counts of neutrophils (Welch’s *t*[14.249] = -4.116, *p* = 0.001). **O** Total T-cell counts (Mann-Whitney *U* = 62, *p* = 0.183). **P** Natural killer cell counts (Student’s *t*[25] = -1.573, *p* = 0.128). **Q** B-cell counts (Mann-Whitney *U* = 66, *p* = 0.251). **R** CD4+ T-cell abundance as a percentage of total T-cells. **S** CD4+ T-cell counts (Mann-Whitney *U* = 81, *p* = 0.683). **T** CD8+ T-cell abundance as a percentage of total T-cells. Data from **R** and **T** were analyzed with a two-way ANOVA due to the interdependence of relative abundance of subtypes (main effect of DSS – *F*[1, 25] = 0.099, *p* = 0.756; main effect of subtype – *F*[1, 25] = 362.027, *p* < 0.001; interaction – *F*[1, 25] = 0.099, *p* = 0.756). **U** CD8+ T-cell counts (Mann-Whitney *U* = 69, *p* = 0.305). We note these counts are extremely low and are likely not reliable. Data in **B-U** are presented as mean ± SEM.\*\**p* < 0.01, \*\*\**p* < 0.001.

## Author’s contributions

JSB, MEK, OUH, and MGT contributed to the overall design and organization of the research. JSB, MEK, JEJ, NKN, CLC, and OUH contributed to the execution of animal studies, including day-to-day animal care and euthanasia. JSB, MEK, and JEJ analyzed data and created figures. JSB, OUH, and MGT wrote and edited the manuscript. JSB performed statistical and bioinformatics analysis. All authors read and approved the final manuscript.

## Acknowledgements

We thank Hannah Staley for her contribution to the multiplexed MesoScale Discovery immunoassays and Karen McFarland, PhD, for her insight into key bioinformatics decisions. We thank LC Sciences for performing the RNA sequencing. We thank the whole Tansey lab for their discussion, insight, and feedback concerning this study. Figures 1A and 3H were created using BioRender.com. Partial funding for this work was derived from awards from the NIH T32-NS082128 (JSB), NIH NIA 1RF1AG057247 (MGT), NIH NINDS 1RF1NS28800 (MGT), and the joint efforts of The Michael J. Fox Foundation for Parkinson’s Research (MJFF) and the Aligning Science Across Parkinson’s (ASAP) initiative. MJFF administers the grant ASAP-020527 on behalf of ASAP and itself. For the purpose of open access, the author has applied a CC-BY public copyright license to the Author Accepted Manuscript (AAM) version arising from this submission.

