## Supplementary figures and images for "A leaky gut dysregulates gene networks in the brain associated with immune activation, oxidative stress, and myelination in a mouse model of colitis"

### Supplemental Figure 1

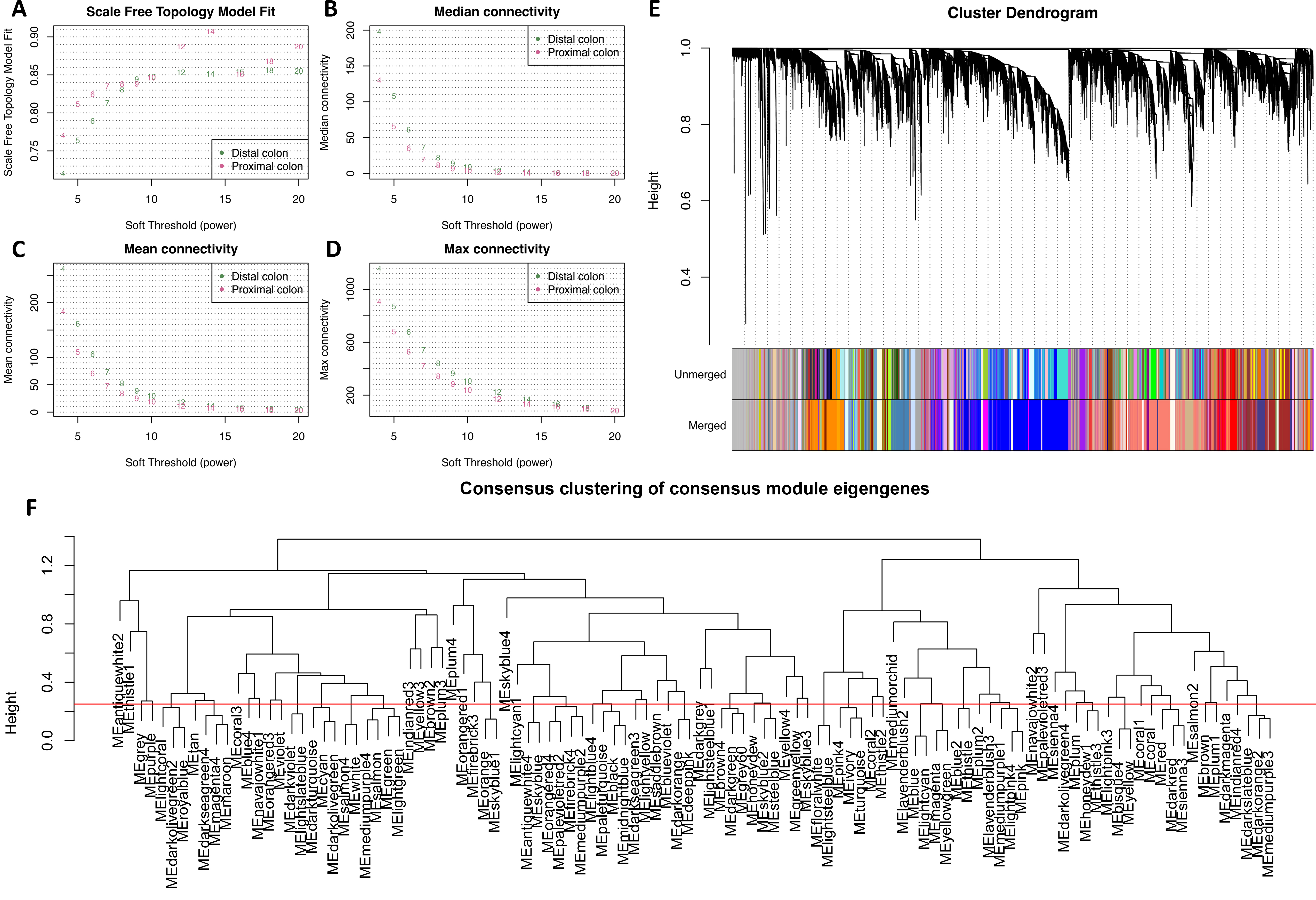

### Supplemental Figure 2

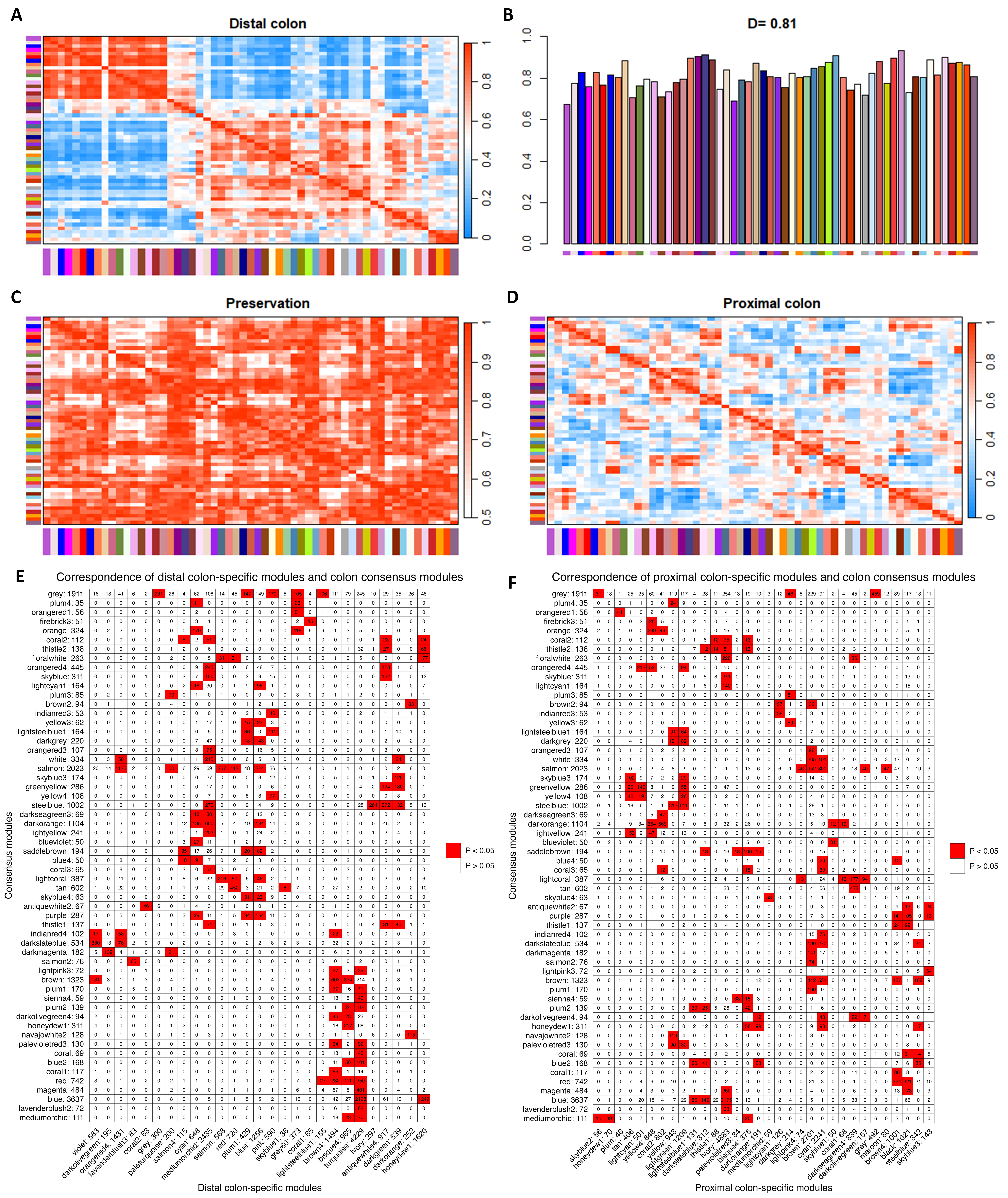

### Supplemental Figure 3

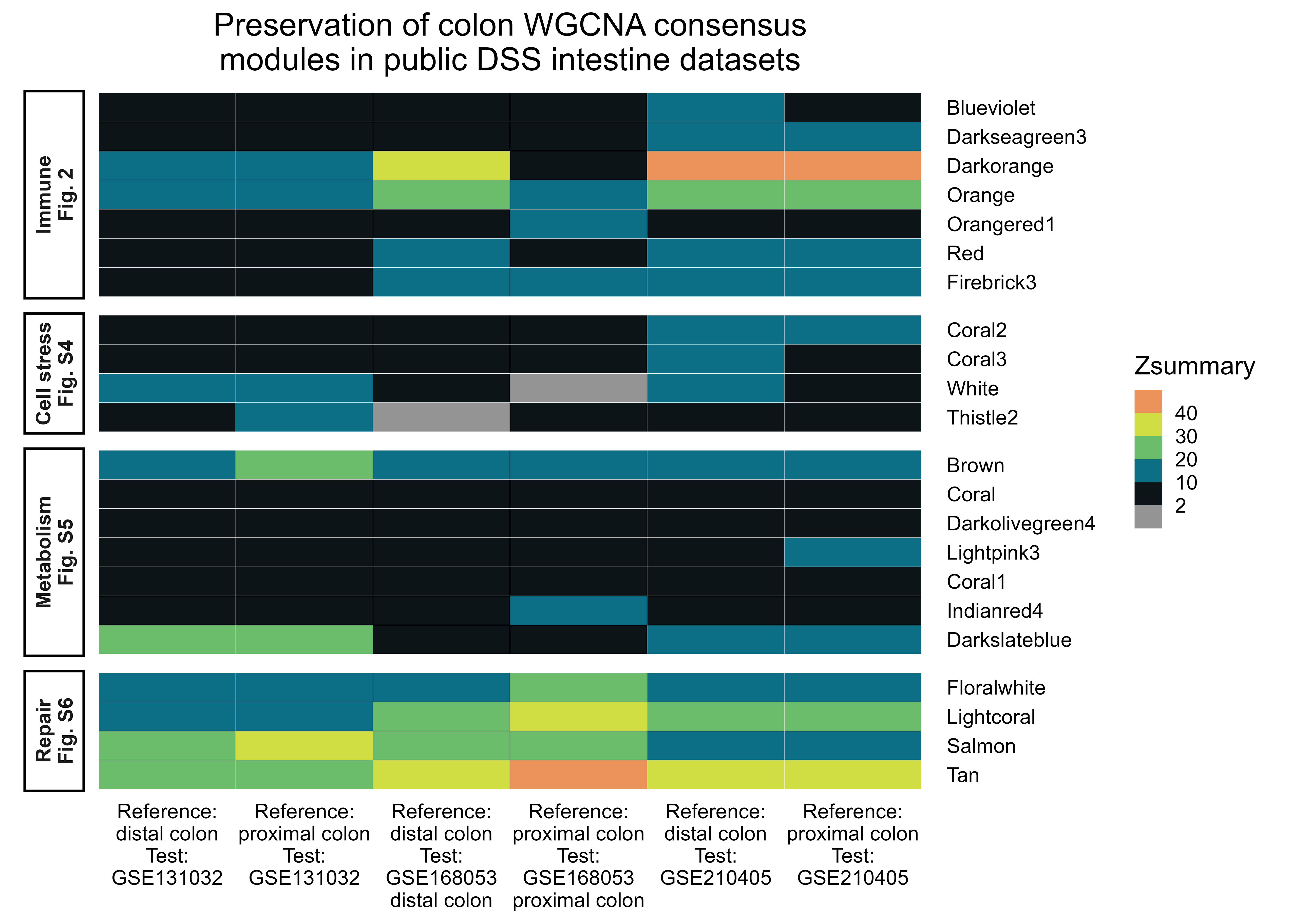

### Supplemental Figure 4

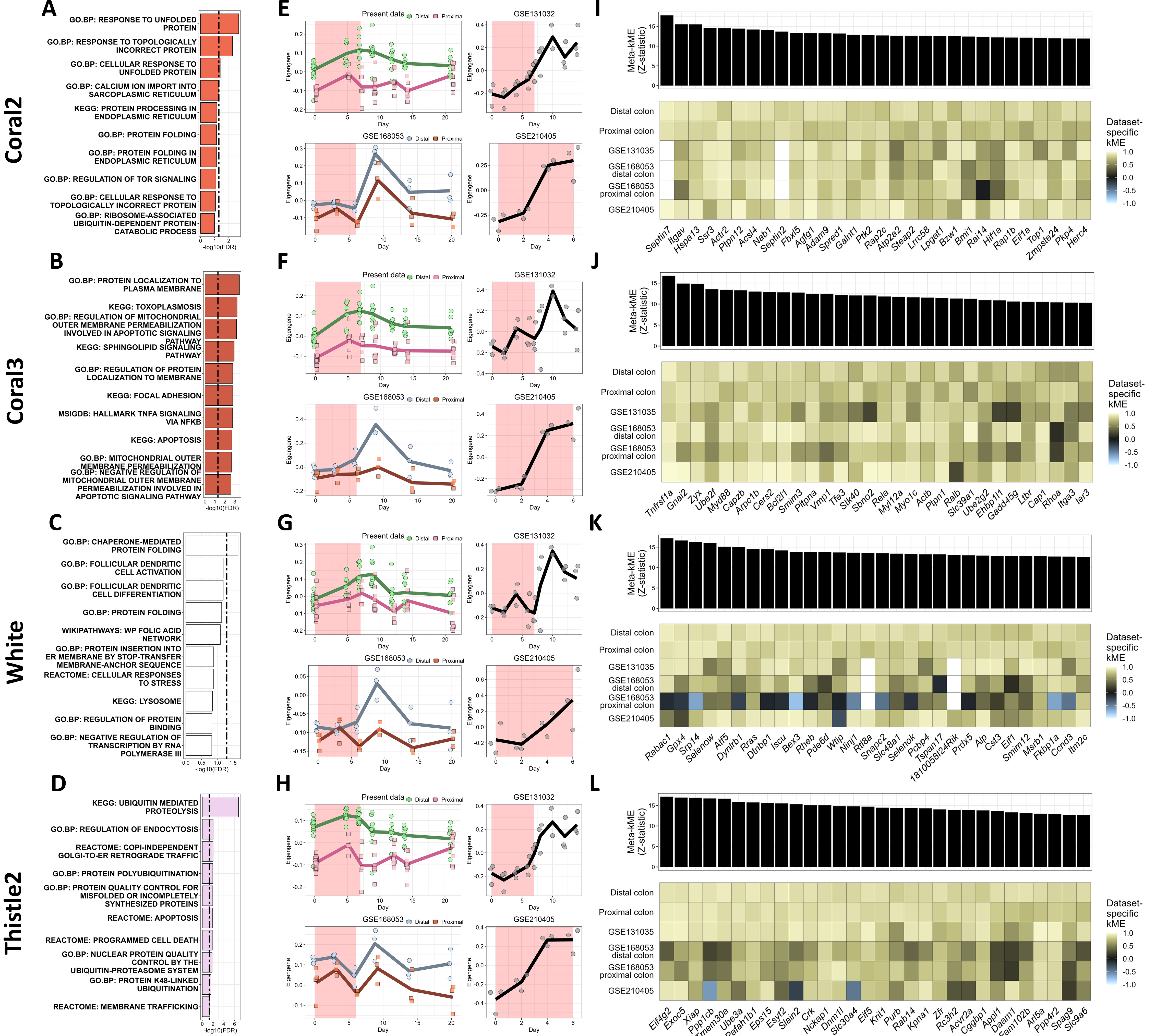

### Supplemental Figure 5

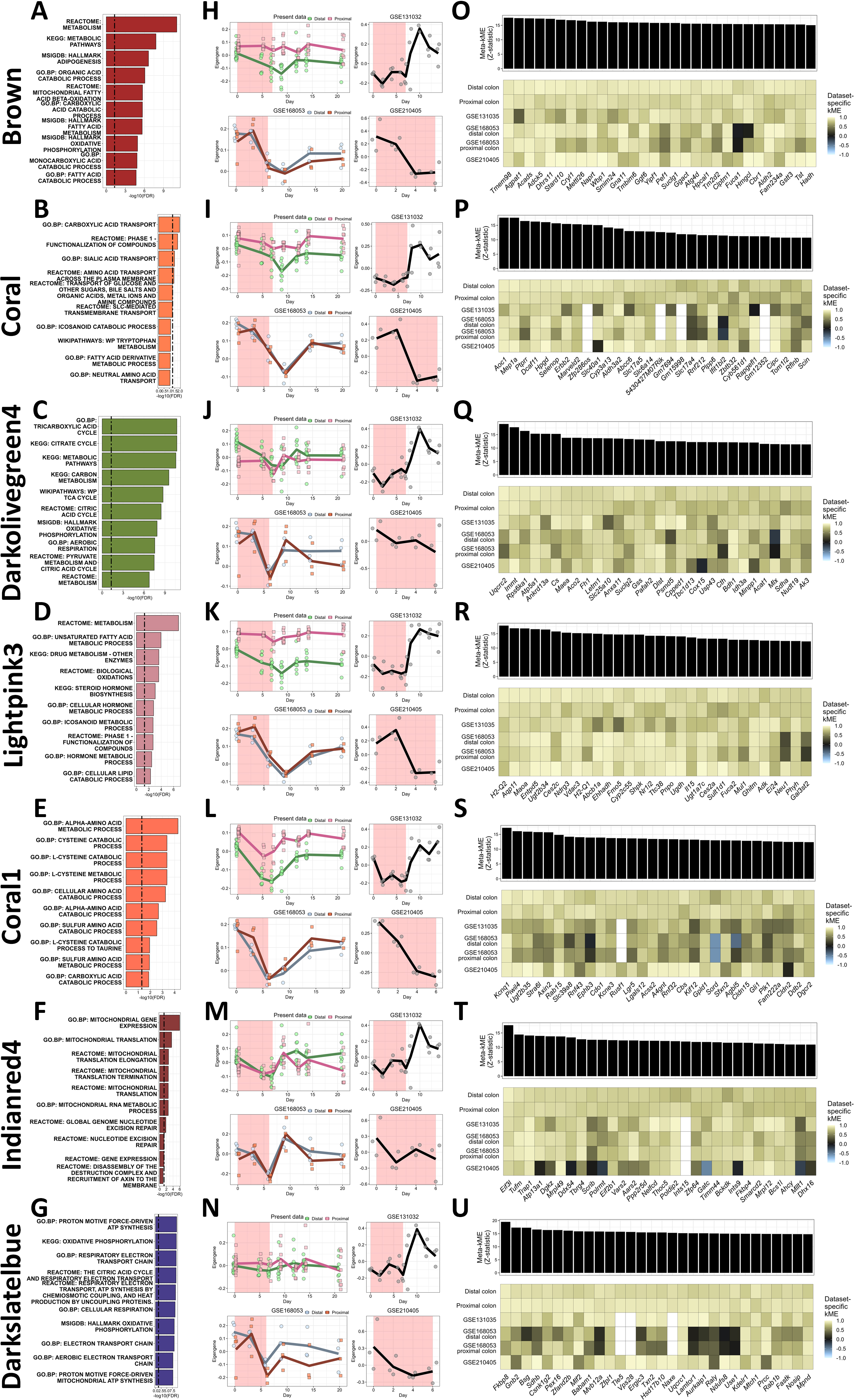

### Supplemental Figure 6

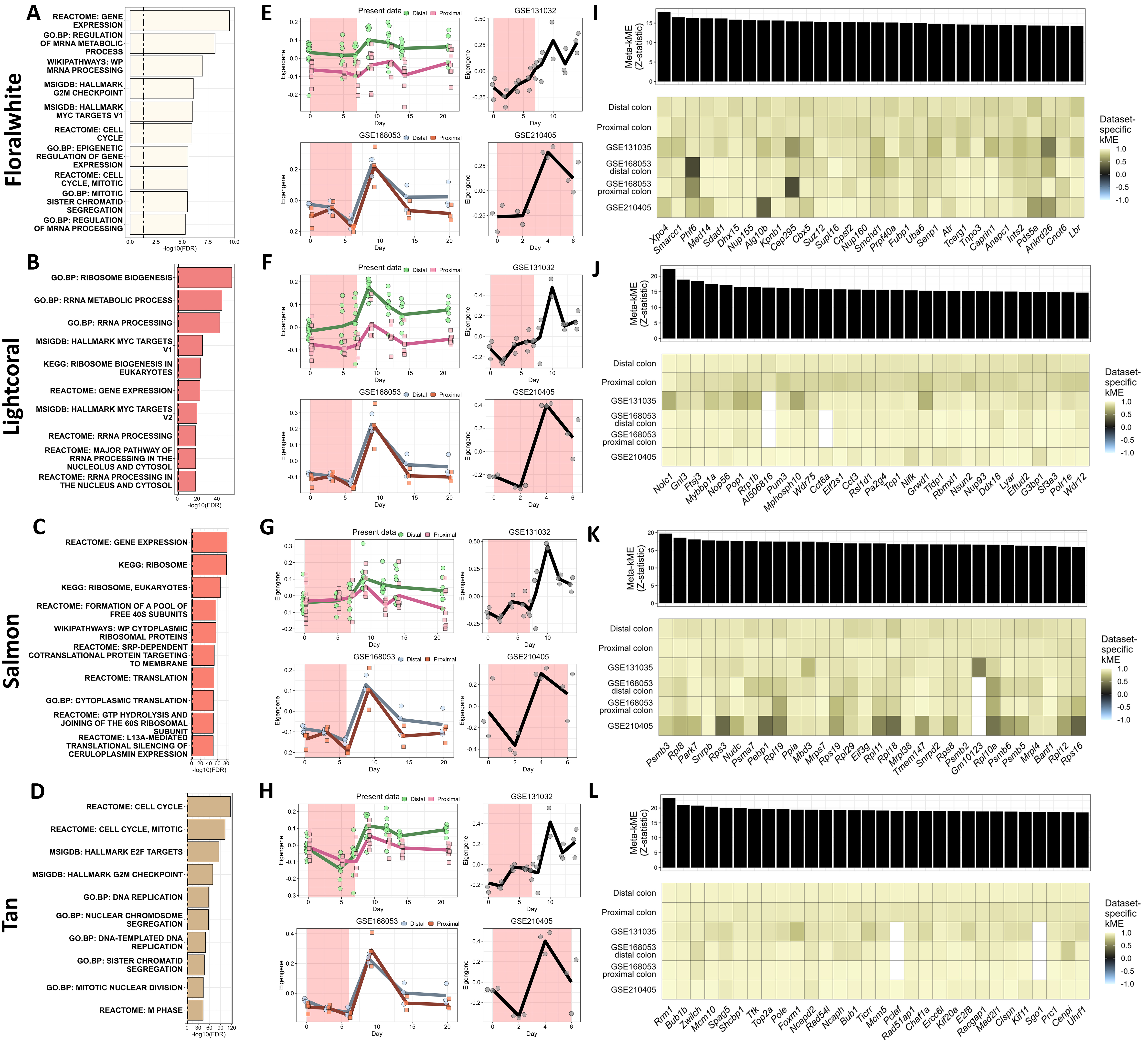

### Supplemental Figure 7

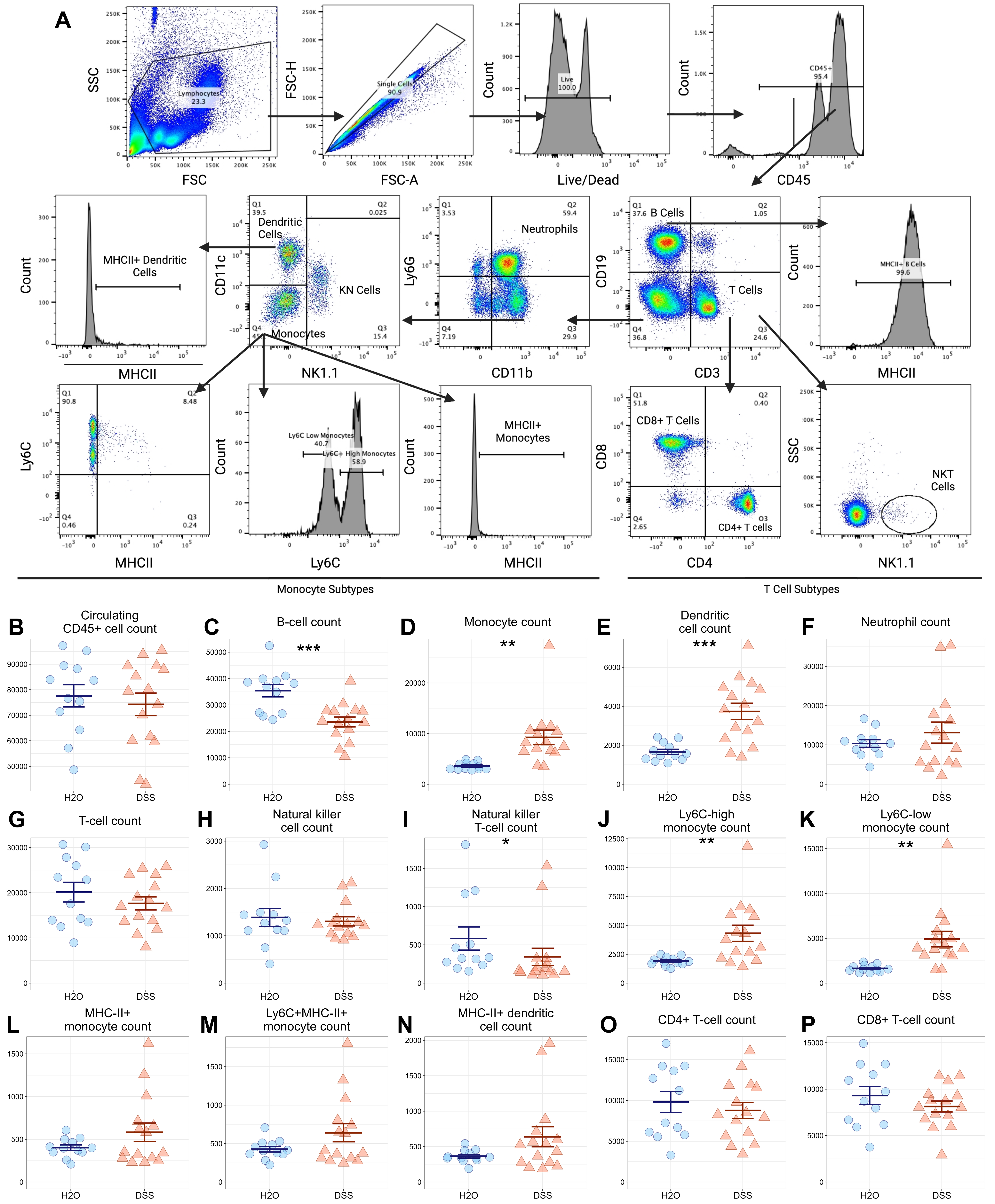

### Supplemental Figure 8

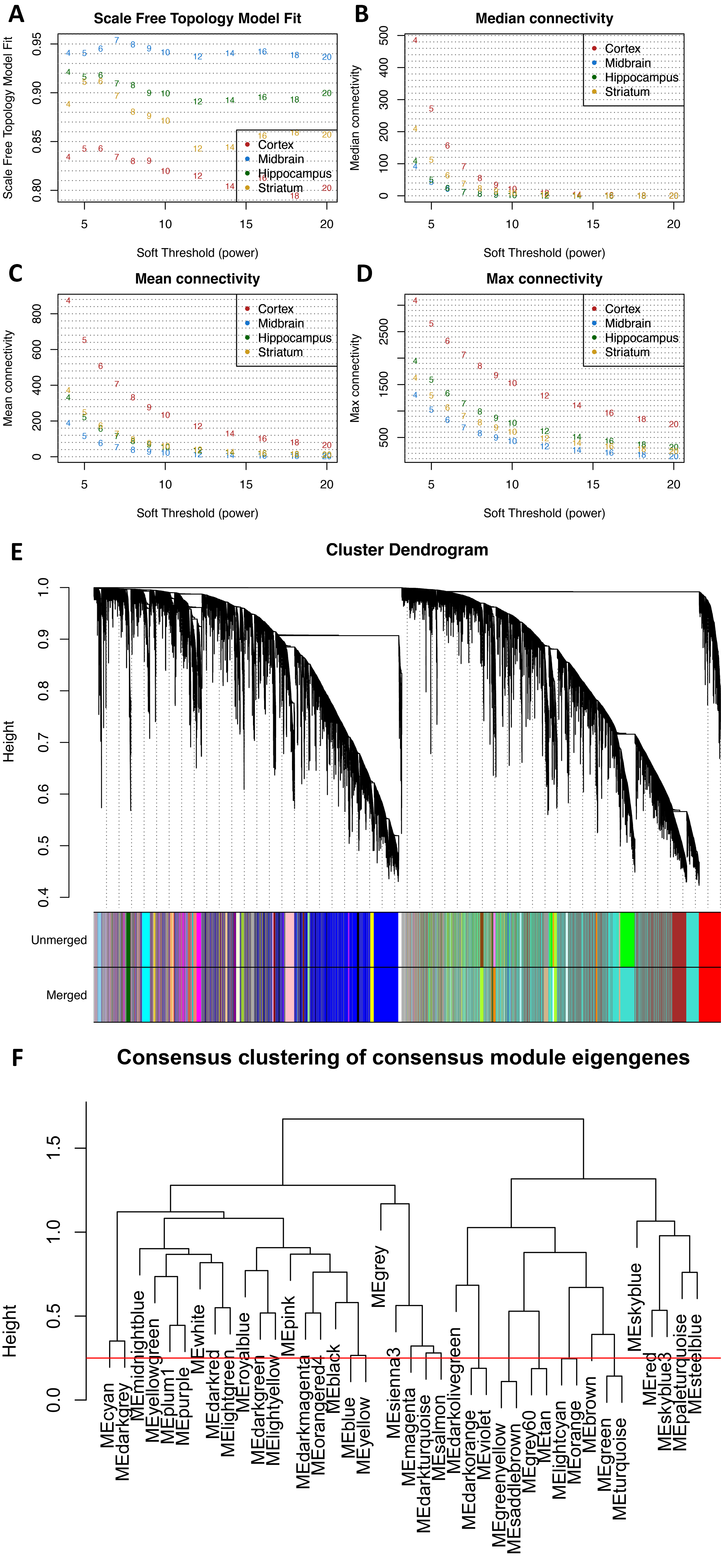

### Supplemental Figure 9

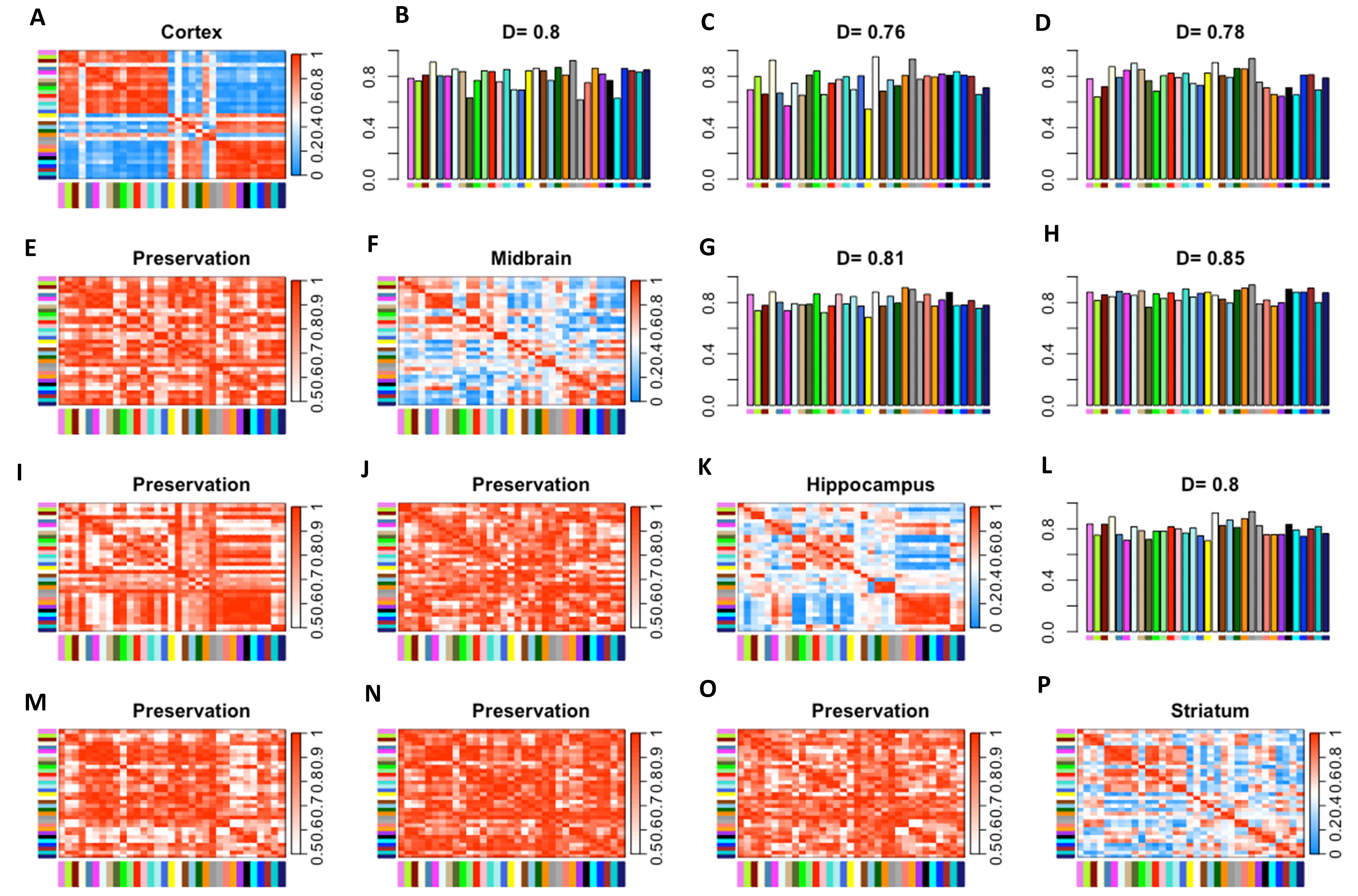

### Supplemental Figure 10

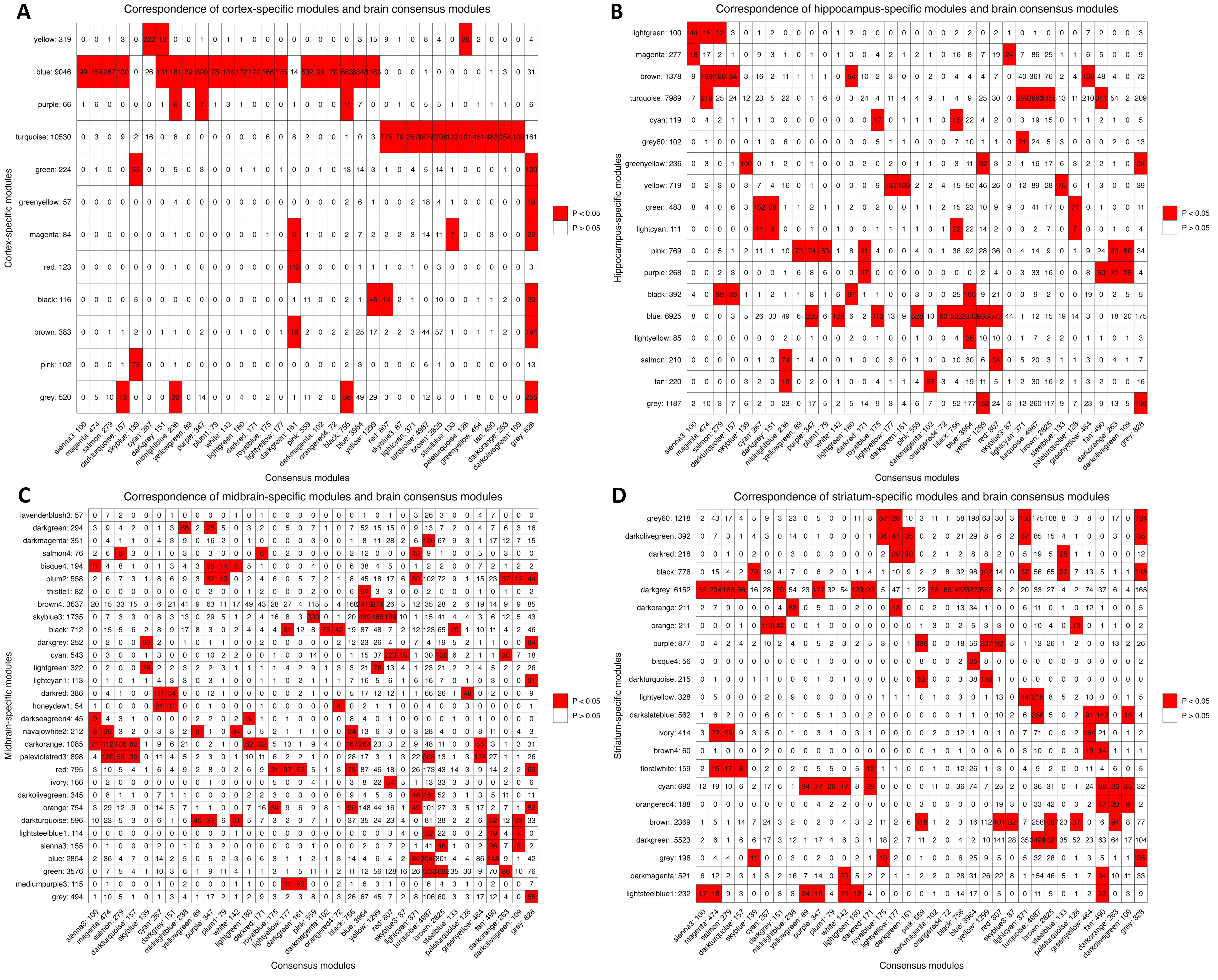

### Supplemental Figure 11

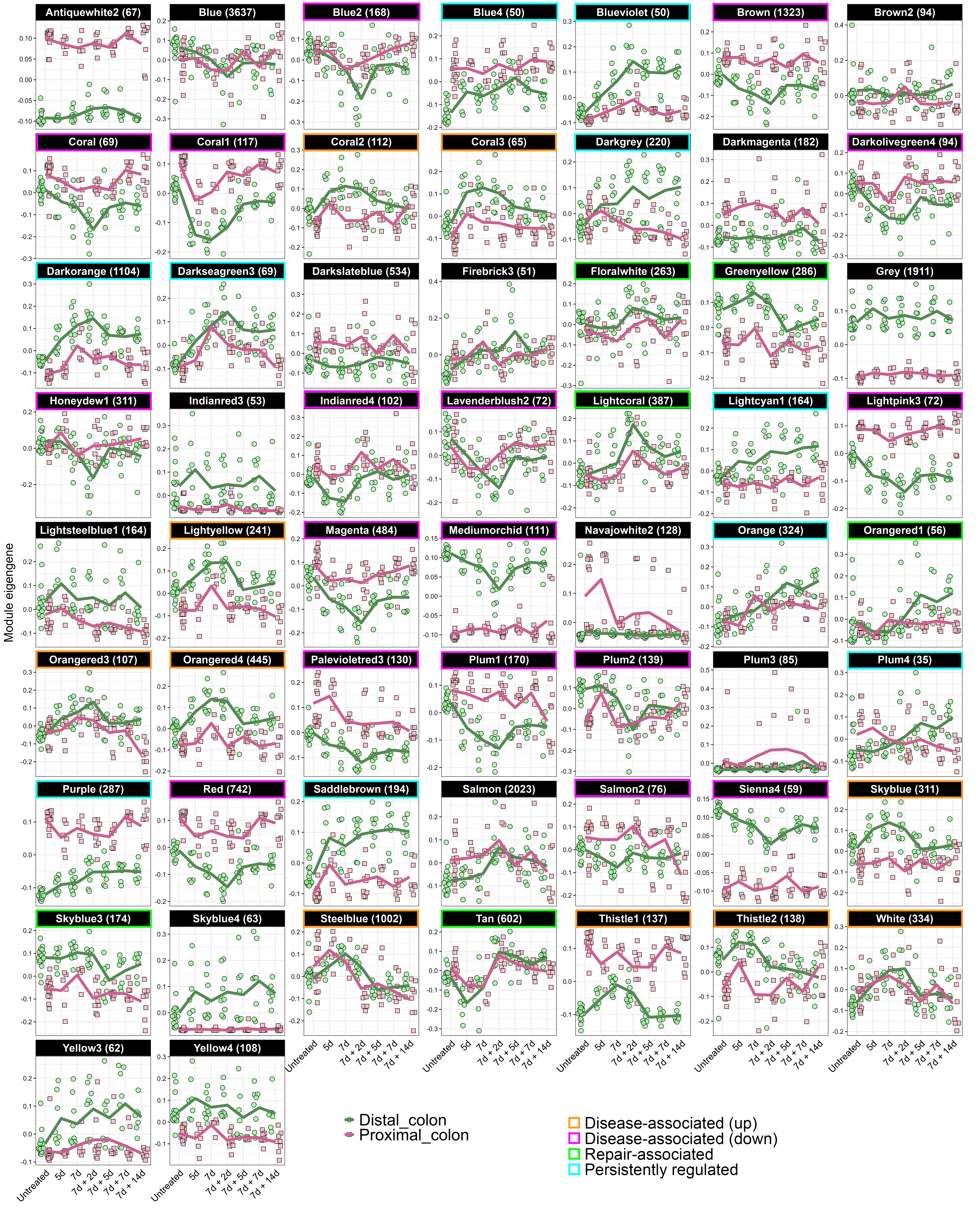

### Supplemental Figure 12

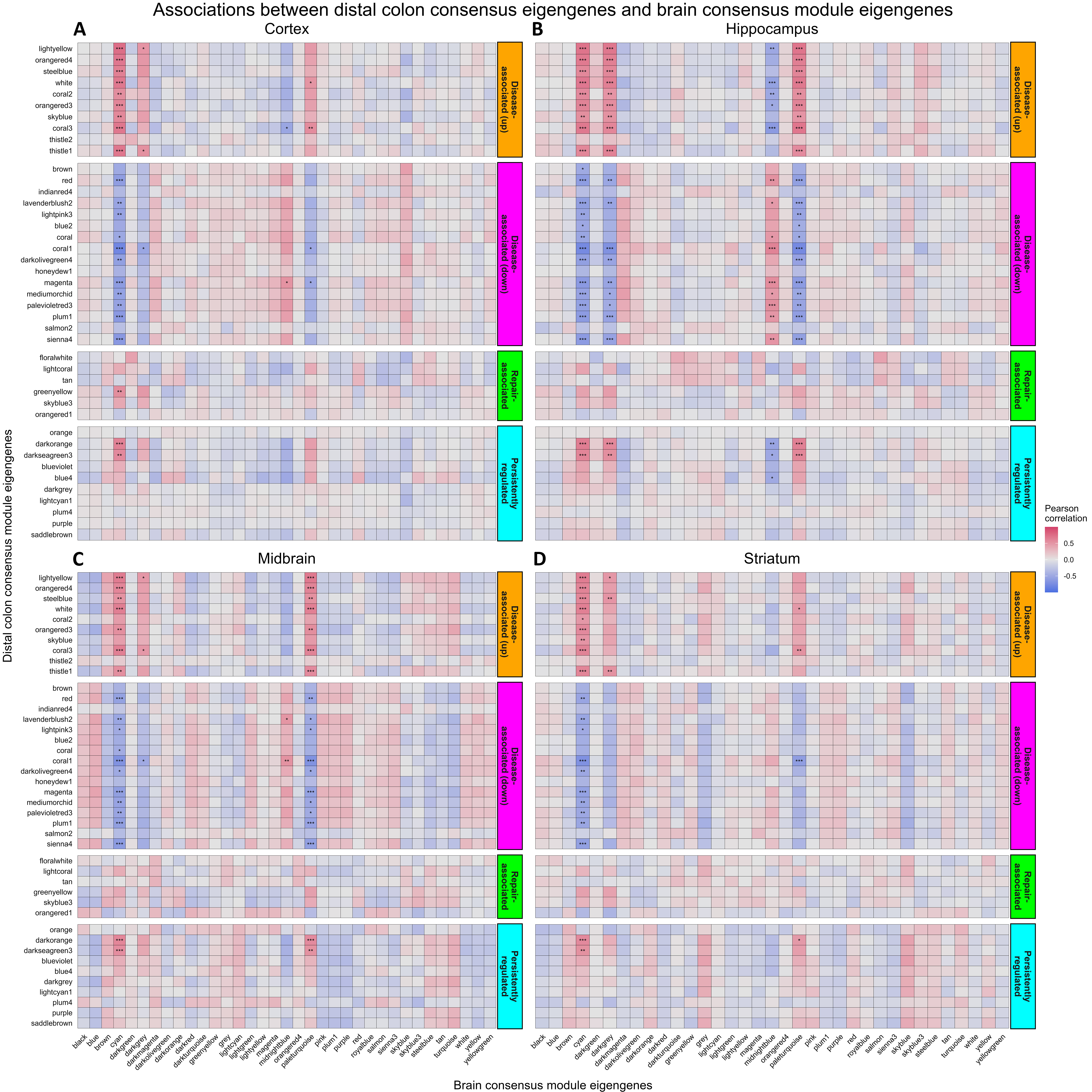

### Supplemental Figure 13

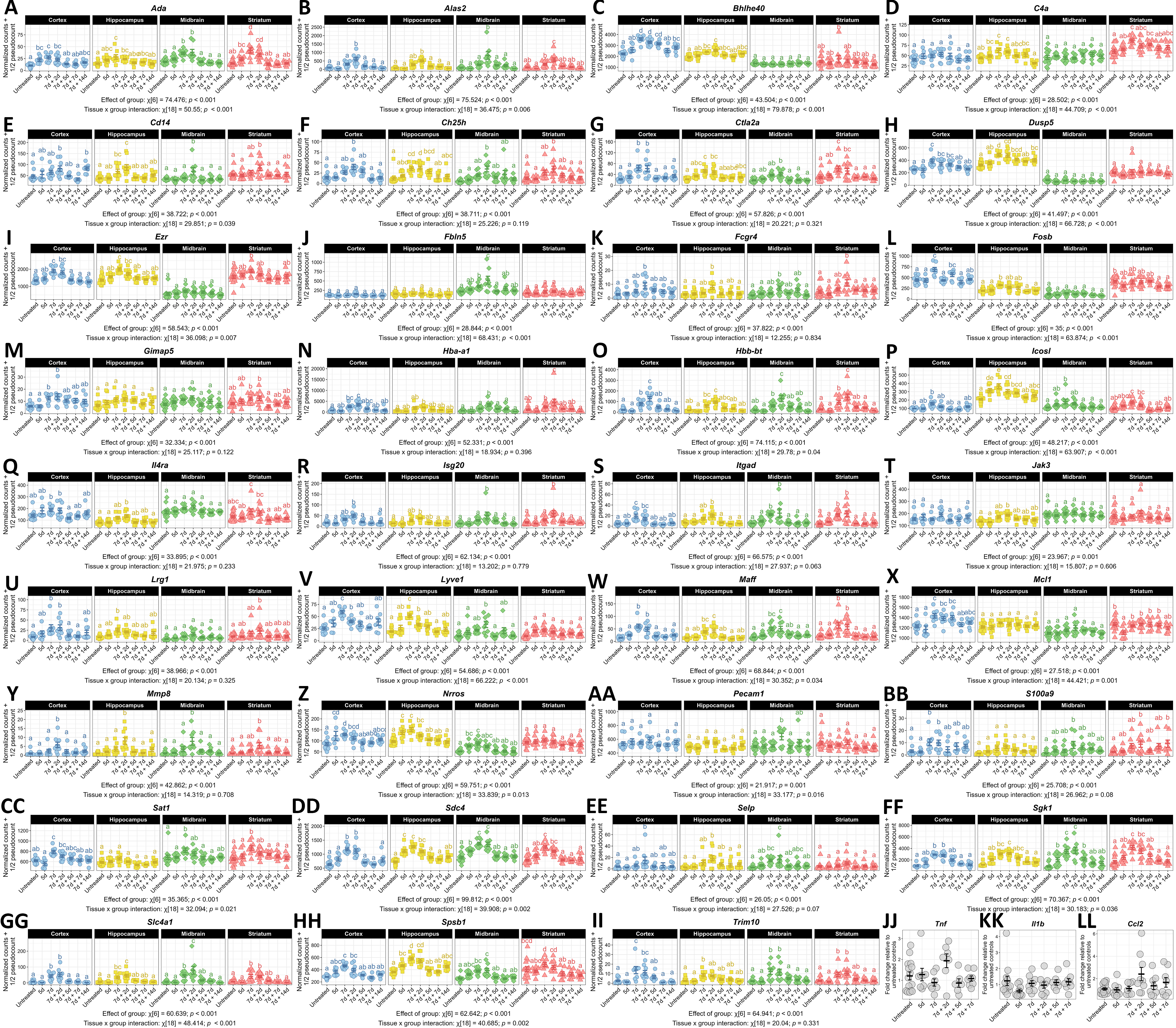

### Supplemental Figure 14

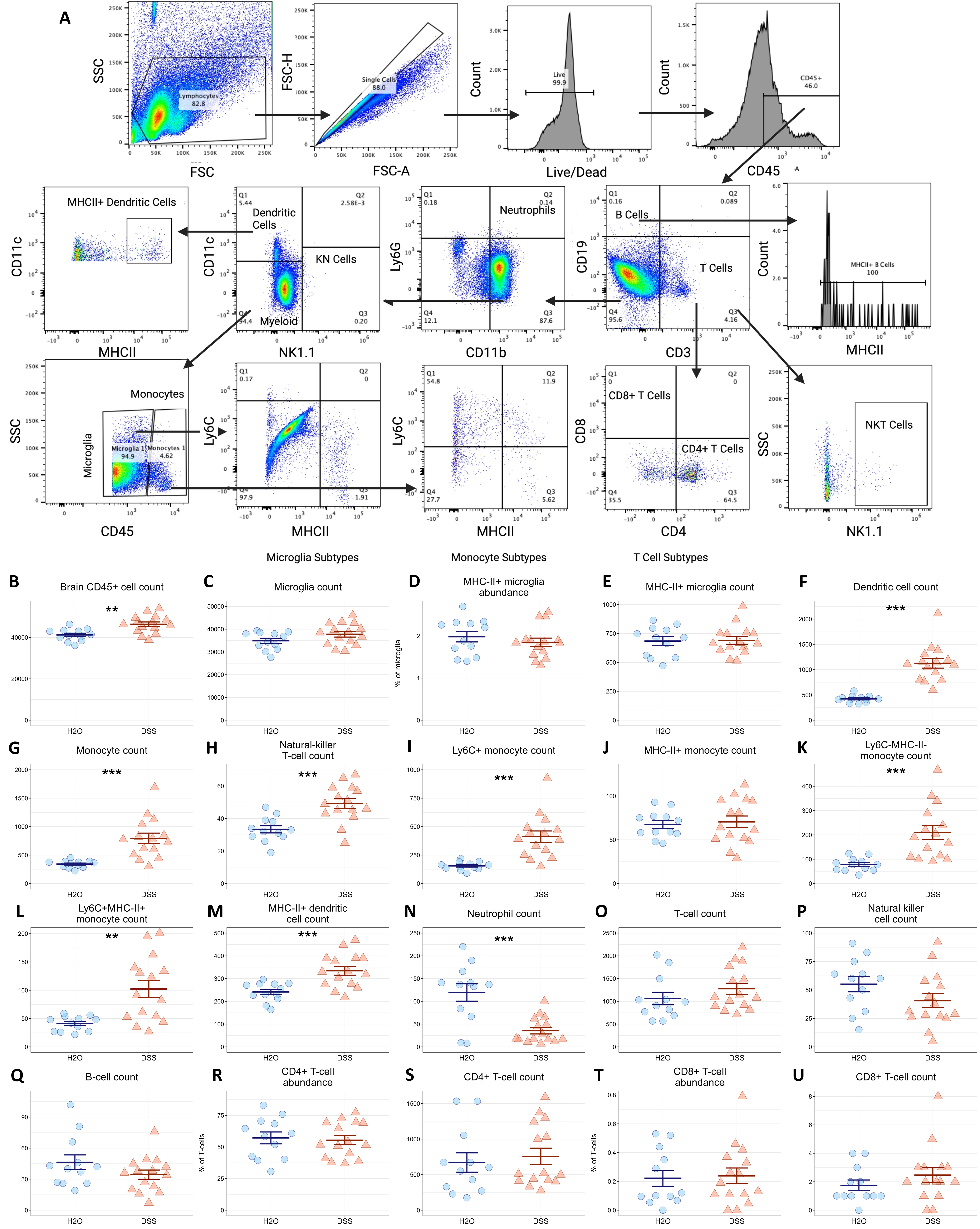
